# Endomembrane systems are reorganized by ORF3a and Membrane (M) of SARS-CoV-2

**DOI:** 10.1101/2021.06.01.446555

**Authors:** Yun-Bin Lee, Minkyo Jung, Jeesoo Kim, Myeong-Gyun Kang, Chulhwan Kwak, Jong-Seo Kim, Ji-Young Mun, Hyun-Woo Rhee

## Abstract

The endomembrane reticulum (ER) is largely reorganized by severe acute respiratory syndrome coronavirus 2 (SARS-CoV-2). SARS-CoV-2 ORF3a and membrane (M) protein expression affects ER-derived structures including cubic membrane and double membrane vesicles in coronavirus-infected cells; however, the molecular mechanisms underlying ER remodeling remain unclear. We introduced a “plug and playable” proximity labeling tool (TurboID-GBP) for interactome mapping of GFP-tagged SARS-CoV-2 ORF3a and M proteins. Through mass spectrometric identification of the biotinylated lysine residue (K+226 Da) on the viral proteins using Spot-TurboID workflow, 117 and 191 proteins were robustly determined as ORF3a and M interactomes, respectively, and many, including RNF5 (E3 ubiquitin ligase), overlap with the mitochondrial-associated membrane (MAM) proteome. RNF5 expression was correlated to ORF3a ubiquitination. MAM formation and secreted proteome profiles were largely affected by ORF3a expression. Thus, SARS-CoV-2 may utilize MAM as a viral assembly site, suggesting novel anti-viral treatment strategies for blocking viral replication in host cells.

**Highlights:**

- SARS-CoV-2 proteins ORF3a and M alter endoplasmic reticulum proteome profile
- ORF3a affects mitochondrial-associated membrane formation
- SARS-CoV-2 may utilize mitochondrial-associated membrane as viral assembly site
- ORF3a and M interactome proteins may serve as targets for COVID-19 treatment

**eTOC Blurb:** ER remodelling by SARS-CoV-2 ORF3a and M protein

## INTRODUCTION

The ongoing pandemic of coronavirus disease 2019 (COVID-19) caused by the severe acute respiratory syndrome coronavirus 2 (SARS-CoV-2) necessitates detailed research into the molecular mechanisms underlying the ability of this virus to infect and colonize host cells. In particular, understanding the mechanism by which each viral protein encoded by the SARS-CoV-2 genome interacts with host factors is essential for the prevention or treatment of COVID-19. Similar to SARS-CoV and Middle East respiratory syndrome coronavirus (MERS), SARS-CoV-2 has been classified in the genus *Betacoronavirus* of the *Coronaviridae* family (Qin et al., 2005). It encodes 16 non-structural proteins (NSP1–16), 4 structural proteins (spike protein [S], membrane protein [M], envelope protein [E], and nucleocapsid protein [N]), and at least 9 accessory proteins (ORF3a, ORF3b, ORF6, ORF7a, ORF7b, ORF8, ORF9b, ORF9c, and ORF10). The non-structural proteins are generated from an auto-proteolytic process and participate in the formation of the replicase-transcriptase complex, which is essential for the viral life cycle (Gordon et al., 2020). The structural proteins are components of each viral particle and are key factors in host cell infection (Gordon et al., 2020). The accessory proteins have functions associated with interferon and immune responses; for example, ORF3a of SARS-CoV activates the NOD-like receptor protein 3 (NLRP3) inflammasome (Siu et al., 2019), and ORF3b of SARS-CoV-2 acts as an interferon antagonist (Konno et al., 2020).

A previous study has demonstrated viral protein–host protein interactions of SARS-CoV-2 using a co-immunoprecipitation (Co-IP) method followed by mass spectrometry analysis (Gordon et al., 2020). The Co-IP can show interaction partners with strong binding affinity to the viral proteins of interest (vPOI); however, these interactome data may also include artificial protein-protein interactions that do not occur in cells since these experiments are conducted using cell lysates isolated in artificial lysis buffer conditions. To overcome these limitations, proximity labeling (PL) methods have recently been developed, which are based on an enzymatic biotinylating reaction using *in situ*-generated reactive biotin species by genetically expressed enzymes. Examples of these enzymes include engineered ascorbate peroxidase (APEX) (Rhee et al., 2013), promiscuous biotin ligase (BioID) (Roux et al., 2012), and TurboID, which is an engineered biotin ligase with faster kinetics (labeling time less than 30 min) (Branon et al., 2018). Since the biotinylation reaction induced by the PL enzymes occurs in living cells, and the labeling radius of the reactive biotin species (biotin-phenoxy radical and biotin-AMP) is less than 50 nm, mass spectrometry analyses of endogenous proteins biotinylated by the PL enzymes can accurately reflect the physiological interactome of the POI that is genetically expressed alongside these enzymes.

PL is being used to identify the host interactome of viral proteins, including coronaviruses (V’Kovski et al., 2019) and SARS-CoV-2 (Laurent et al., 2020; Samavarchi-Tehrani et al., 2020; St-Germain et al., 2020). Although this method shows various important host proteins that support viral life cycles (such as eIF3 and eIF4 complexes) and anti-viral proteins (such as MAVS, PKR, and LSM14A/B), the results from this technique may contain false positive findings due to the conventional mass detection workflow. Currently, most PL studies (including the viral interactome studies) have used conventional mass detection of non-biotinylated peptides of the streptavidin-bead (SA-bead) enriched proteins from the PL-labeled cell lysate. However, this workflow can misidentify artificial interacting proteins in the lysis buffer as biotinylated proteins (false positive) when those proteins are reproducibly shown in the biotinylated samples over the controls. Furthermore, recent reports describing the mapping of the SARS-CoV-2 interactome (Laurent et al., 2020; St-Germain et al., 2020) utilized BioID, which performs slow biotinylating reactions on proximal proteins over a 12-h period. This long biotinylation reaction may perturb the physiological conditions of host cells.

To maximize the advantages of recently improved PL methods, in this study, we combined TurboID, a fast biotinylating enzyme (Branon et al., 2018), with the Spot-BioID method (Lee et al., 2019). In this technique, cell lysates are digested with trypsin followed by SA-bead enrichment at the peptide level, so that only peptides modified by TurboID with biotin-conjugated lysine (K+226) are detected using mass spectrometry. We further assessed the ability of our modified PL method, dubbed *Spot-TurboID*, to analyze proteins biotinylated by TurboID, and to ultimately obtain *in vivo* host interactome information without false positive results.

In the present study, we aimed to identify which proteins located near the endoplasmic reticulum (ER) interact with SARS-CoV-2 proteins, as the ER membrane is proposed to be the primary replication site for SARS-CoV such as SARS-CoV-2 (Stertz et al., 2007). Folding and synthesis of membrane, lipids, secretory proteins, sterols, and ion storage occurs in the ER (Lin et al., 2008). Among SARS-CoV-2 proteins, ORF3a is associated with vesicle sorting and fusion mechanisms while the M protein interacts with membrane-shaping proteins (St-Germain et al., 2020). Therefore, we focused on the ORF3a and M proteins, using our modified PL system to examine their functions in living cells. We conducted correlative light and electron microscopy (CLEM) imaging experiments to determine the localization and targeting sites of these proteins in living cells. These findings can offer new research directions to improve our understanding of the mechanisms underlying SARS-CoV-2 infection and to identify new therapeutic targets for COVID-19.

## Results

### GFP-tagged SARS-CoV-2 structural and accessory proteins target specific organelles

To confirm the subcellular localization of the SARS-CoV-2 proteins, we prepared 11 constructs of vPOIs (NSP7, NSP8, NSP9, M, N, ORF3a, ORF3b, ORF6, ORF7b, ORF9b, and ORF9c) tagged with GFP using a 13-amino acid linker (GAPGSAGSAAGSG) (**Figure 1A**). Confocal imaging of HEK293-AD cells showed that the GFP-tagged vPOIs were well expressed (**Figure 1B**). Compared with non-structural proteins, structural and accessory proteins targeted specific organelles. NSP7, NSP8, and NSP9 showed a whole-cell expression pattern, indicating that these proteins were diffused throughout the cell, and do not specifically interact with a certain organelle, possibly due to the lack of viral RNAs. In contrast, M, N, ORF3a, ORF3b, ORF6, ORF7b, ORF9b, and ORF9c showed organelle-specific localization patterns, indicating that these proteins interact with host cell proteins of specific organelles. Among these constructs, we observed that the expression patterns of ORF3a and M merged well with an ER marker protein, SEC61B (**Figure S1A**).

**Figure 1.**
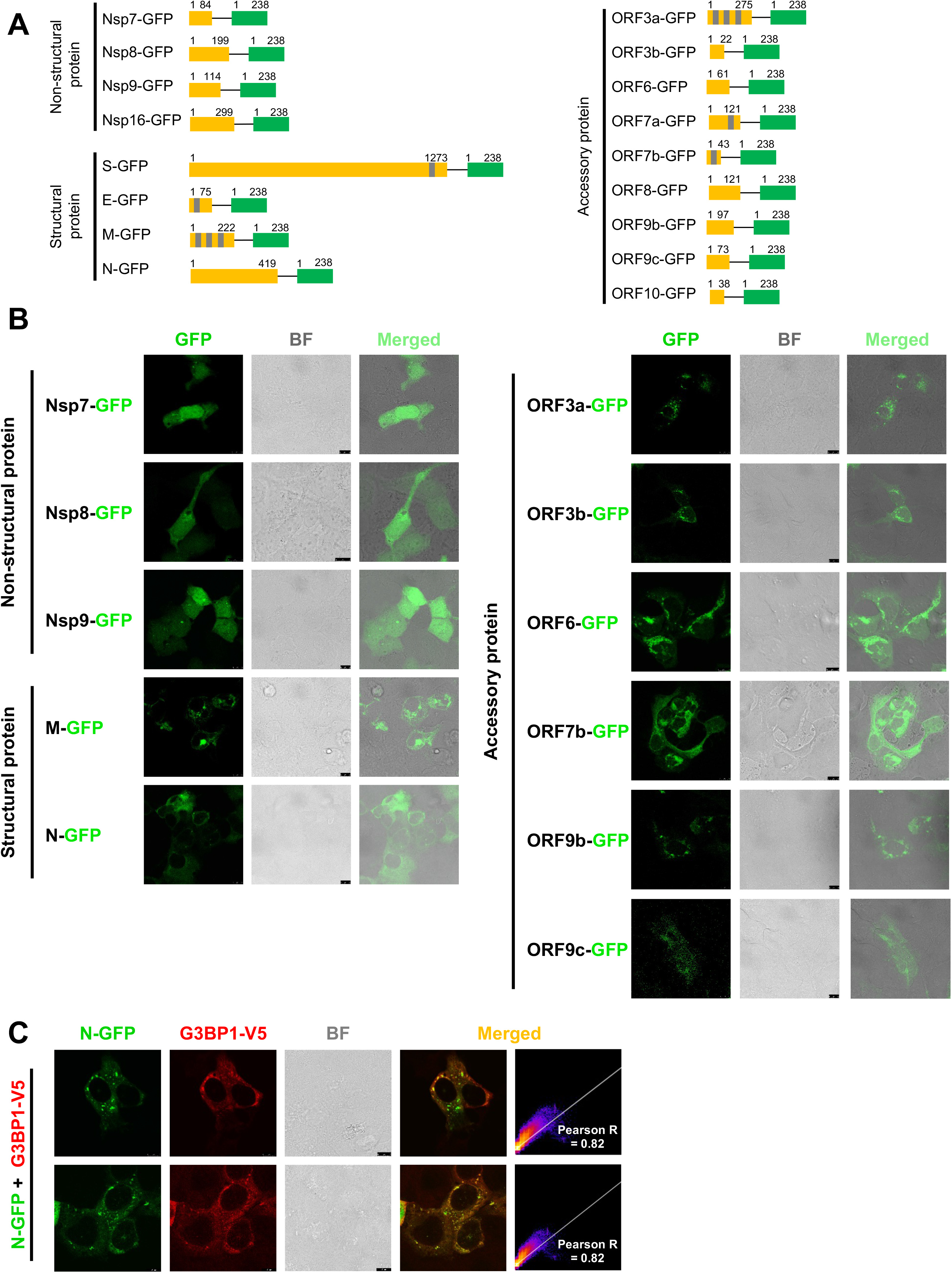
Confocal imaging of GFP-tagged vPOI of SARS-CoV-2. **(a)** Schematic view of viral protein of interest (vPOI)-linker-GFP constructs (linker sequence: GAPGSAGSAAGSG). Predicted transmembrane (TM) domains by TMHMM program are shown in light gray. **(b)** Confocal microscopy results of vPOI-GFP constructs in HEK293-AD cells. Scale bars: 10 μm. **(c)** Confocal microscopy images of co-expressed N-GFP and G3BP1-V5. G3BP1-V5 was visualized using anti-V5 antibody (AF568-conjugated, red fluorescence channel). Pearson correlation values were calculated between different fluorescent channel images. Scale bars: 8 μm (upper) and 10 μm (below). BF: bright field.

In line with a previous study showing that N protein is associated with stress granules (Gordon et al., 2020), confocal imaging of HEK293-AD cells co-expressing N-GFP and G3BP1-V5 (stress granule marker protein) (Anderson and Kedersha, 2006; Nover et al., 1983), showed that the two patterns clearly merged, representing a stress granule pattern (**Figure 1C**). In addition, we confirmed that GFP-tagged vPOIs (i.e. ORF3a, ORF6, ORF7, M) were merged with Flag-tagged vPOIs in co-expressed cells, indicating that tagging with GFP (~27 kDa) is acceptable for those vPOIs like as tagged with small epitope tag (Flag) (**Figure S1B)**. These experiments confirmed that the GFP-tagged structural and accessory protein constructs of SARS-CoV-2 established were suitable for our further interactome study.

### The GFP/GBP binding system is applicable for identifying the SARS-CoV-2 interactome in live cells

As the POI-GFP and APEX2-GFP binding protein (GBP) system was previously validated for GFP-specific electron microscope (EM) imaging (Ariotti et al., 2015), we utilized this GFP and GBP binding “plug and play” system for the interactome analysis of vPOI-GFP by co-expression of TurboID-GBP via *Spot-TurboID* workflow (**Figure 2A**). To confirm a specific binding between vPOI-GFP and TurboID-GBP, M, N, ORF3a, ORF6, and ORF7b were selected. Confocal images of HEK293-AD cells co-expressing vPOI-GFP and TurboID-GBP were identical to those of cells expressing vPOI-GFP alone (**Figure 2B, S2**), demonstrating that GFP and GBP bound well, and the patterns of the biotinylated proteins merged with the GFP and GBP patterns. Thus, we could confirm that co-expressed TurboID-GBP has a strong and specific binding interaction with GFP-tagged vPOIs and it did not appear to interrupt the targeting of vPOI-GFP to specific organelles.

**Figure 2.**
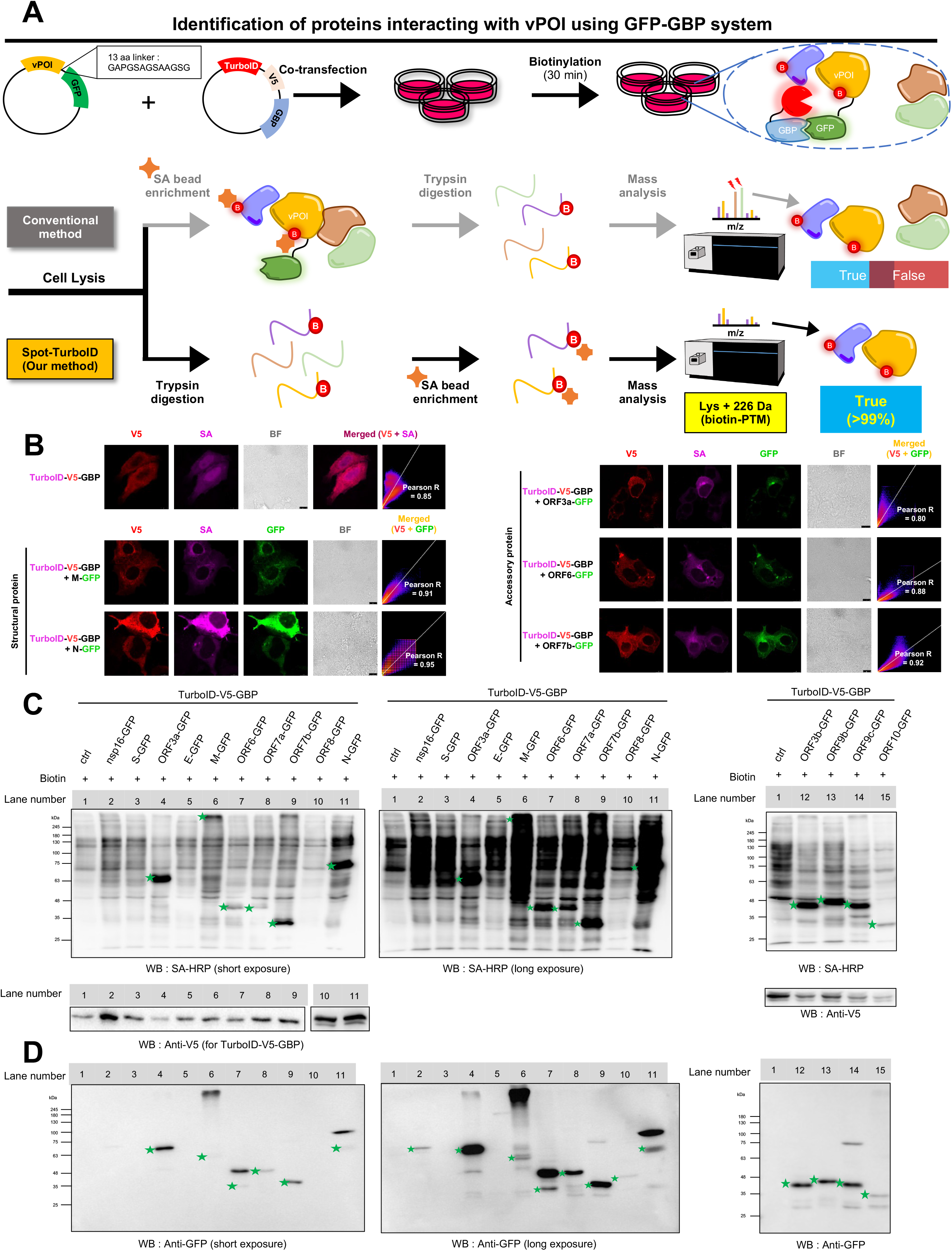
Interactome mapping of SARS-CoV-2 viral proteins by *Spot-TurboID* workflow. **(a)** Overview of our *Spot-TurboID* method combined with vPOI-GFP and TurboID-GBP binding system **(b)** Confocal microscopy images of vPOI-GFP and TurboID-V5-GBP in HEK293-AD cells. TurboID-V5-GBP was visualized by anti-V5 antibody (AF568-conjugated, RFP fluorescence channel). Biotinylated proteins were visualized by AF647-conjugated streptavidin (Cy5 fluorescence channel). Scale bars: 10 μm. Additional images are shown in **Figure S2**. BF: bright field. **(c)** Streptavidin-horseradish peroxidase (SA-HRP) western blotting results of biotinylated proteins by TurboID-GBP under the co-expression of vPOI-GFP in HEK293T cells. Band signals of biotinylated vPOI-GFP by TurboID-GBP are marked with green asterisks. Anti-V5 blot signals of the same samples are shown below. **(d)** Anti-GFP western blotting results of the same samples as **(c)**. Bands of the expected molecular weight of each vPOI-GFP constructs are marked with green asterisks. Raw images of blot results are shown in **Figure S3**.

In the western blot analysis of proteins biotinylated by TurboID-GBP and detected using streptavidin-horseradish peroxidase (SA-HRP), we observed that the biotinylated protein patterns of TurboID-GBP were altered when it was co-expressed with GFP-tagged M, N, ORF3a, ORF3b, ORF6, ORF7a, ORF7b, ORF9b, ORF9c, or ORF10 proteins (**Figure 2C, D, S3A-C**). The change in SA-HRP western blot patterns of the biotinylated proteins by the PL enzyme (Lee et al., 2016) may reflect that the neighboring proteins around TurboID-GBP were significantly altered when TurboID-GBP was translocated to the co-expressed vPOI-GFP in the same cell. These experiments also showed that the GFP-tagged structural and accessory proteins of SARS-CoV-2 may be surrounded by different host proteins in cells, and that these proteins may be readily biotinylated by TurboID and identified using mass spectrometry via the *Spot-TurboID* workflow (**Figure 2A**).

### ORF3a and M SARS-CoV-2 proteins destructively perturb the ER membrane organization

Among the structural and accessory proteins that exhibit an ER targeting pattern, ORF3a and M contain 3 transmembrane (TM) domains (**Figure 3A, B, S4A-B**) and both proteins showed a similar alteration of protein band patterns in western blot analysis, with a bigger molecular weight than expected (**Figure 2D**). ORF3a shows 85.1% sequence similarity with ORF3a of SARS-CoV (Gordon et al., 2020), which exhibits ion transport activity and induces NLRP3 inflammasome activation (Siu et al., 2019). ORF3a has also been shown to possess pro-apoptotic activity (Ren et al., 2020). Similarly, M of SARS-CoV-2 shows 96.4% sequence similarity with M of SARS-CoV (Gordon et al., 2020), which was reported to target the ER, ER-Golgi intermediate compartment, and Golgi apparatus, and is associated with apoptosis (Chan et al., 2007) and nuclear factor-(NF) κB signaling (Fang et al., 2007). We hypothesized that these proteins may perturb the ultrastructure of endomembrane systems, as it has been shown that many viral proteins target the endomembrane systems of host cells (Inoue and Tsai, 2013).

**Figure 3.**
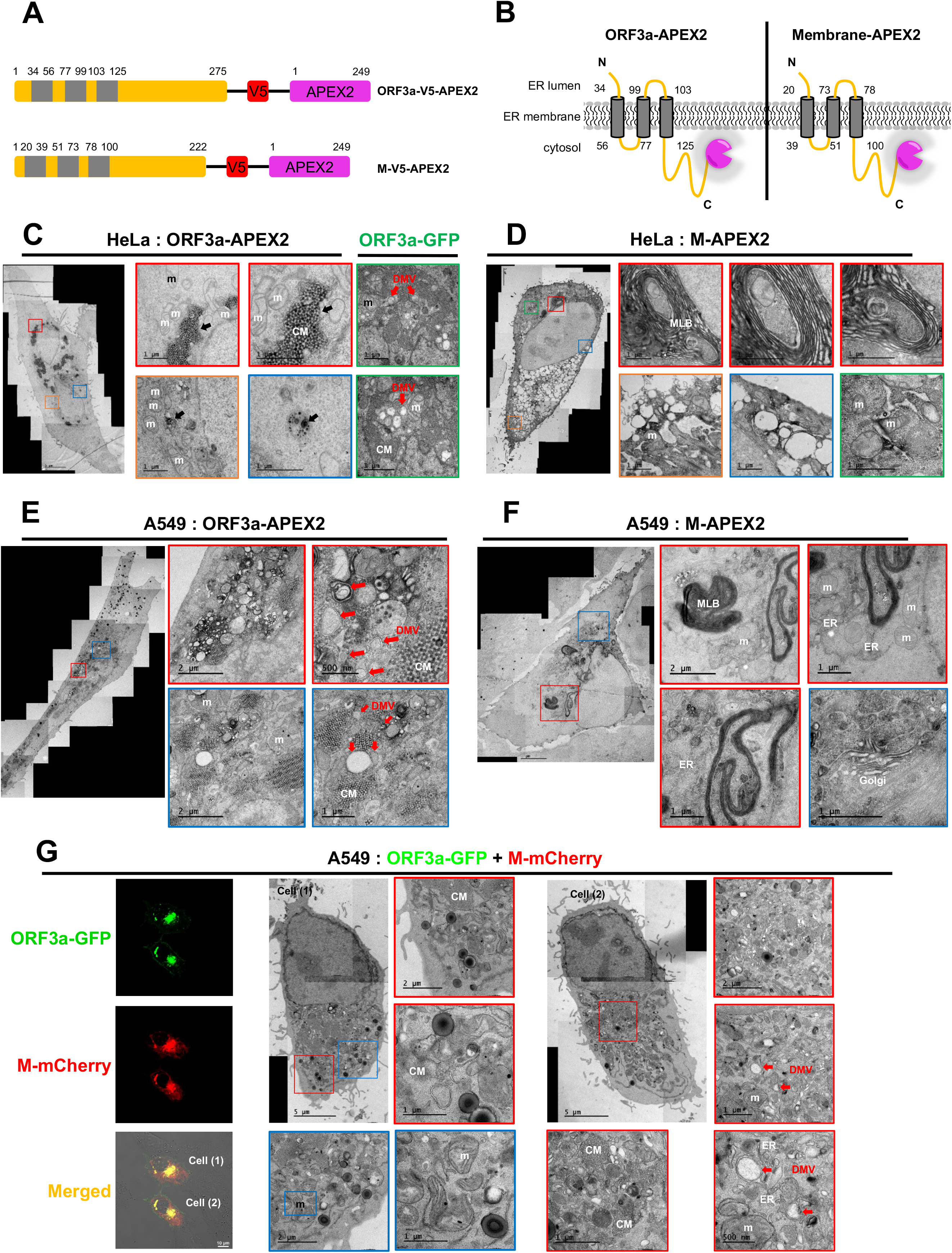
EM results of ORF3a and Membrane protein of SARS-CoV-2 expressed cells. **(a)** Schematic view of ORF3a-linker-V5-APEX2 and M-linker-V5-APEX2 constructs. Linker sequence: GAPGSAGSAAGSG. TM domains of vPOI are shown in light gray. **(b)** Proposed membrane topology of ORF3a-APEX2 and M-APEX2 at the ER membrane **(c)** EM images of DAB-stained ORF3a-APEX2- and ORF3a-GFP-transfected HeLa cells. Scale bars: 1 μm. Distinct membrane structures were observed with higher magnification. Magnified areas are marked with different colored boxes. Additional EM images are shown in **Figure S5a.** Parts with red arrows mark DMV structures (c, e, g). Mitochondria and cubic membrane are indicated as “m” and “CM”, respectively (c–g). **(d)** EM images of M-APEX2 transfected HeLa cells. Scale bars: 1 μm. Additional EM images are shown in **Figure S5b. (e)** EM images of ORF3a-APEX2 transfected A549 cells. Scale bas: 500 nm–2 μm. **(f)** EM images of M-APEX2 transfected A549 cells. Scale bars: 1 μm– 2 μm. **(g)** CLEM images of ORF3a-GFP and M-mCherry co-transfected A549 cells. Ultrastructures of endomembrane systems in each cell were observed by EM imaging. Scale bars: 500 nm–5 μm.

To test this hypothesis, we prepared ORF3a-APEX2 and M-APEX2 constructs, with APEX2 tagged to the C-terminus of the vPOIs (**Figure 3A**), and used them for EM imaging in HeLa cells following APEX-mediated diaminobenzidine (DAB) staining (Martell et al., 2012). ORF3a and M proteins largely disrupted the ER **(Figure 3C, D; Figure S5A-B)**. ORF3a-APEX2 expression significantly increased cubic membrane (CM, also denoted as convoluted membrane) structure formation; moreover, these CM structures were generated in ORF3a-GFP expressing cells shown by CLEM imaging (**Figure 3C, S5C**). Notably, CM structures also form in coronavirus-infected cells (Deng and Angelova, 2021; Snijder et al., 2020), and they are considered as neo-organelles for the assembly of viral proteins (Uchil and Satchidanandam, 2003). Additionally, in ORF3a-GFP expressed HeLa cells, structure of double membrane vesicle (DMV, red arrows in **Figure 3C**) was discovered near ER and mitochondria. Formation of DMV by ORF3a-APEX2 was reproduced in A549 lung cell line (**Figure 3E, G).** We confirmed that these CM and DMV structures were not observed in SEC61B-APEX2 or APEX2 transfected controls (**Figure S6A-C**). Since DMV formation is believed to house replication complexes of various RNA viruses (Cortese et al., 2020; Klein et al., 2020; Wolff et al., 2020), our result indicates that ORF3a of SARS-CoV-2 plays an important role in the formation of ER-derived neo-organelles (i.e. CM and DMV) that are essential for viral replication in host cells.

Similarly, ER was clearly disrupted in M-expressing cells (**Figure 3D**) and appeared to curl into whorl patterns (also termed as aggresomes). These patterns were also observed in herpes simplex virus-infected cells (Nii et al., 1968) and ER stress-induced cells (Schuck et al., 2014; Snapp et al., 2003). In CLEM imaging of M-GFP (**Figure S5D**), we observed the formation of multilamellar bodies (MLB) at the ER membrane (**Figure 3D, F**) and electron-dense autophagic vesicles (green arrows in **Figure S5D**). The biogenesis of MLB regulated is by autophagy (Hariri et al., 2000) and MLBs are utilized during extracellular vesicle-type viral egress (Bunyavirus) (Sanz-Sánchez and Risco, 2013). In flavivirus replication (such as Zika virus), autophagy is usually activated and autophagic vesicles accumulate inside the cells in *in cellulo* (Hamel et al., 2015) and *in vivo* (Cao et al., 2017) experiments. In our study, these structures were not detected in non-transfected HeLa (**Figure S5B**) and A549 cell controls **(Figure S6A**). Moreover, we confirmed that the structure of ER, Golgi, and mitochondria did not change in SEC61B-APEX2 or APEX2-expressed cells (**Figure S6B-C**). Our EM results therefore imply that the ORF3a and M proteins of SARS-CoV-2 may largely contribute to the generation of ER-derived neo-organelles (Fernández de Castro et al., 2014; Tenorio et al., 2018; Xie et al., 2019) for efficient viral assembly and replication by remodeling the endomembrane systems of host cells. To identify the host proteins involved in this process, we conducted an *in cellulo* interactome analysis of the SARS-CoV-2 ORF3a and M proteins using GFP-tagged constructs with TurboID-GBP via the *Spot-TurboID* workflow.

### ORF3a and M interactomes showed perturbed proteomic landscape at the ER membrane

For mass spectrometric analysis, we used the aforementioned *Spot-TurboID* method (**Figure 2A**) to obtain an accurate interactome map of the ORF3a and M proteins. To obtain the biotinylated peptidome, HEK293T cells co-expressing vPOI-GFP and TurboID-GBP and control HEK293T cells co-expressing GFP and TurboID-GBP (**Figure 4A**) were treated with biotin (50 μM, 30 min) in full 10% fetal bovine serum medium, washed with cold Dulbecco’s phosphate-buffered saline, and lysed using 2% sodium dodecyl sulfate in 1× Tris-buffered saline. Lysate samples were trypsin-digested prior to SA-bead enrichment for biotinylated peptidome enrichment. Biotinylated peptides were isolated from the SA-beads and injected to the Orbitrap Fusion Lumos mass spectrometer. A total of 4232, 3818, and 6299 peptide-spectrum matches containing biotin-moiety peptides at the lysine residue (K+226) were detected in the ORF3a-GFP + TurboID-GBP, M-GFP + TurboID-GBP, and unconjugated GFP (control) + TurboID-GBP expressing cells, respectively.

**Figure 4.**
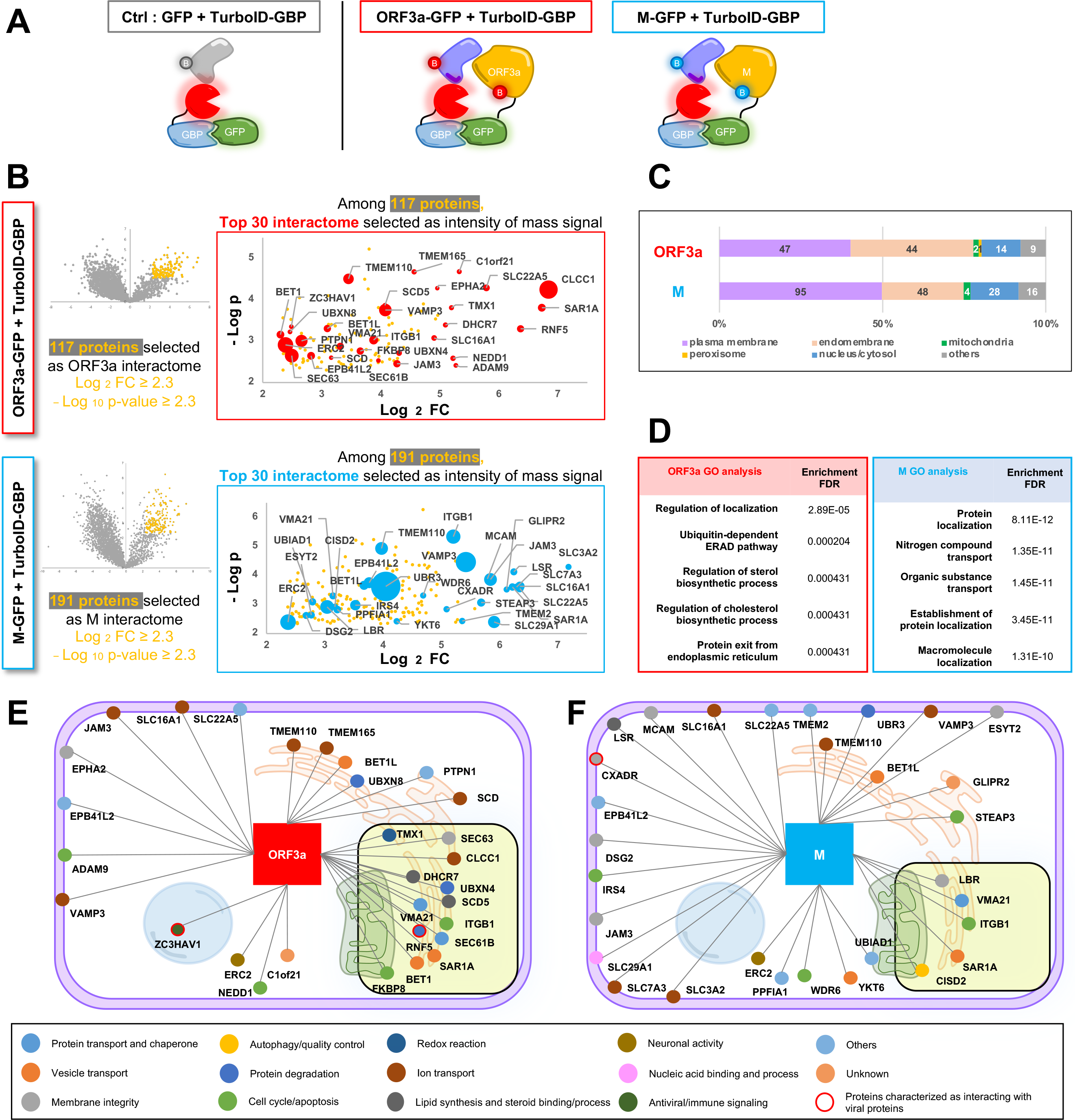
Mass analysis of host interactome of ORF3a and M protein of SARS-CoV-2 by *Spot-TurboID*. **(a)** Schematic view of three groups of sample for mass analysis: TurboID-GBP + GFP (left, control), TurboID-GBP + ORF3a-GFP (middle), and TurboID-GBP + M-GFP (right). **(b)** Enrichment of the *Spot-TurboID* detected biotinylated proteins by TurboID-GBP in the samples of ORF3a-GFP (upper) or M-GFP (below) over GFP (control), respectively. x-axis: Log2 value of fold change (FC) of mass signal intensities. cut off value is 2.3; y-axis: -Log10 value of p-value, cut off value is 2.3. Gene names of the top 30 proteins are shown in the graph. Bubble size indicated the mass signal intensity. See detailed information in **Figure S7e** and **Dataset S1**. **(c)** Subcellular distribution of the selected interactome of ORF3a (117 proteins) and M (191 proteins). See detailed information in **Figure S7f** and **Dataset S1**. **(d)** Gene ontology analysis (http://bioinformatics.sdstate.edu/go/) of enriched functions of ORF3a and M’s interactomes. See detailed information in **Figure S7g**. **(e, f)** Subcellular map of top 30 selected interactome of ORF3a (e) and M (f). MAM proteins are shown in yellow-colored box.

Based on the reproducibility and similarity values of the biotinylated peptidome of each sample (Lee et al., 2016), replicate samples under the same conditions were clustered together, confirming the reproducibility of the experiment (similarity value > 0.84, **Figure S7A-D**). From the volcano plot analysis of mass signal intensity of proteins biotinylated by TurboID-GBP, proteins that were more strongly biotinylated in the ORF3a-GFP and M-GFP co-expressed samples versus the control GFP-expressed sample were selected as interactome proteins of ORF3a and M, respectively (log2(fold change, FC) **≥**2.3 and –log10 p-value **≥**2.3, **Figure 4B, S7E**). The major population of the interactomes for ORF3a and M comprises endomembrane system proteins, suggesting that SARS-CoV-2 ORF3a and M are primarily localized at the ER.

Among the 117 filtered proteins for the ORF3a interactome, 40.2% were located at the plasma membrane and 37.6% were located at the endomembrane (**Figure 4C, S7F**). Among the 191 filtered proteins of the M interactome, 49.7% were located at the plasma membrane and 25.1% were located at the endomembrane (**Figure 4C, S7F**). Thus, a large proportion of plasma membrane and endomembrane proteins were present in both, the ORF3a and M interactomes. According to gene ontology analysis, proteins associated with regulation of localization and ubiquitin-dependent ER-associated protein degradation (ERAD) pathway were highly enriched among the 117 proteins of the ORF3a interactome (**Figure 4D, S7G**). Proteins related to localization and nitrogen compound transport were highly enriched among the 191 proteins comprising the M interactome (**Figure 4D, S7G**). From these analyses, we could see that the molecular functions of ORF3a and M are highly related to localization and transport of other substances within the endomembrane of host cells.

To further our understanding of these interactions, we then filtered the top 30 most strongly biotinylated proteins (red or blue-colored dots in **Figure 4B**) among the ORF3a and M interactomes, respectively. The 30 filtered proteins of the ORF3a interactome were classified according to localization and function (**Figure 4E**). Surprisingly, 13 of these 30 proteins (43.3%) overlapped with the recently identified mitochondrial-associated membrane (MAM) proteome determined using Contact-ID (Kwak et al., 2020). This suggests that ORF3a of SARS-CoV-2 is localized at the MAM, in line with a previous host interactome analysis of SARS-CoV-2 using the Co-IP method (Davies et al., 2020). Since the MAM-enriched ORF3a interactome proteins are associated with vesicle transport (BET1, BET1L, SAR1A, and VAMP3), regulation of membrane integrity (EPHA2, JAM3, and SEC63), and lipid synthesis and steroid binding/processes (DHCR7 and SCD5), we postulated that ORF3a may be involved in the regulation of vesicle trafficking at the MAM to support viral assembly and egress processes similar to other viral proteins that target the MAM, such as the NS5A and NS5B proteins in hepatitis C virus (Hamamoto et al., 2005). Moreover, several ion transport proteins that are part of the endomembrane system (such as CLCC1, SCD, SLC16A1, SLC22A5, TMEM110, and TMEM165) were identified in the ORF3a interactome. Based on previous findings that ORF3a of SARS-CoV regulates the ion channels at the ER membrane (Chan et al., 2009; Issa et al., 2020) and ORF3a of SARS-CoV-2 forms Ca^2+^ and K^+^ channels (Kern et al., 2020), we postulate that ORF3a may localize at the sub-domain of ER where ion transport proteins cluster.

Similarly, the top 30 filtered proteins of the M interactome were classified by localization and molecular function (**Figure 4F**). Among these proteins, we found the proteins associated with cell cycle/apoptosis (ITGB1, IRS4, WDR6, and STEAP3) to be highly enriched in our M interactome. We also observed that several vesicle transport proteins (BET1L, SAR1A, VAMP3, and YKT6) were highly enriched in the M interactome, which may influence ER vacuolization and apoptosis associated with the M protein (Chan et al., 2007) (**Figure 3D, F**).

Moreover, several known viral interacting host proteins, such as CXADR and ZC3HAV1, were found in the M interactome. Interestingly, infection of SARS-CoV-2 induced an increase in the transcript level of CXADR (coxsackievirus and adenovirus receptor) (Dörner et al., 2004; Martino et al., 2000) in a SARS-CoV-2 infected patient sample (Zhou et al., 2020). Among our M interactome proteins, ZC3HAV1 (CCCH-type zinc-finger antiviral protein) inhibits viral replication by recruiting other proteins responsible for sensing numerous viral RNAs (Kerns et al., 2008; Oshiumi et al., 2015), including human immunodeficiency virus- (HIV)-1 (Zhu et al., 2011), Moloney and murine leukemia virus (Lee et al., 2013), and xenotropic MuLV-related virus (Wang et al., 2012b). Notably, ZC3HAV1 was also enriched in the ORF3a interactome. Thus, our data suggest that ZC3HAV1 is also involved in the anti-viral response against SARS-CoV-2.

### Spot-TurboID reveals topology of TM proteins interacting with ORF3a and M

Since ORF3a and M are targeted to ER membrane (**Figure S1A**), it is reasonable that majority of their interacting proteins are membrane residents such as endomembrane proteins or plasma membrane proteins, synthesized at the ER membrane: ORF3a: 78%, M: 75% **(Figure 4C**). For membrane proteins, identification of their membrane topologies is crucial to understanding their domain-wise function at either side of the membrane. In our work, membrane topological information of the biotin-labeled membrane proteins was easily extracted. Using our *Spot-TurboID* method, we obtained mass spectrometry data that included the biotinylation sites of each digested peptides. All digested peptides isolated following SA-bead enrichment had at least one biotin-modified site at a lysine (K) residue. Since the biotinylation by TurboID-GBP occurred via cytosolic regions of the protein, biotinylated sites on proximal interacting proteins of vPOI-GFP logically faced the cytosol. We then obtained membrane topological information for proteins harboring a TM domain using mass spectrometry data. It is noteworthy that using similar approaches, we successfully revealed the topologies of inner mitochondrial membrane (Lee et al., 2016) and MAM proteins (Kwak et al., 2020).

Using this workflow, 22 TM proteins were identified in each of the ORF3a and M interactomes and the biotinylation sites and topology of these proteins are shown in the figures (**Figure 5, S8A-B**). For example, the biotinylation sites of SEC61B and VAMP3 (K20 of SEC61B, K35 of VAMP3) from the ORF3a interactome are well matched to previously characterized topologies. Based on these results confirming the reliability of the *Spot-TurboID* data, we propose the hitherto unknown topologies of multiple ORF3a and M interacting membrane proteins (CLCC1, DHCR7, TMEM110, FKBP8, RNF5, UBR3, and UBIAD1) (yellow-colored proteins in **Figure 5**).

**Figure 5.**
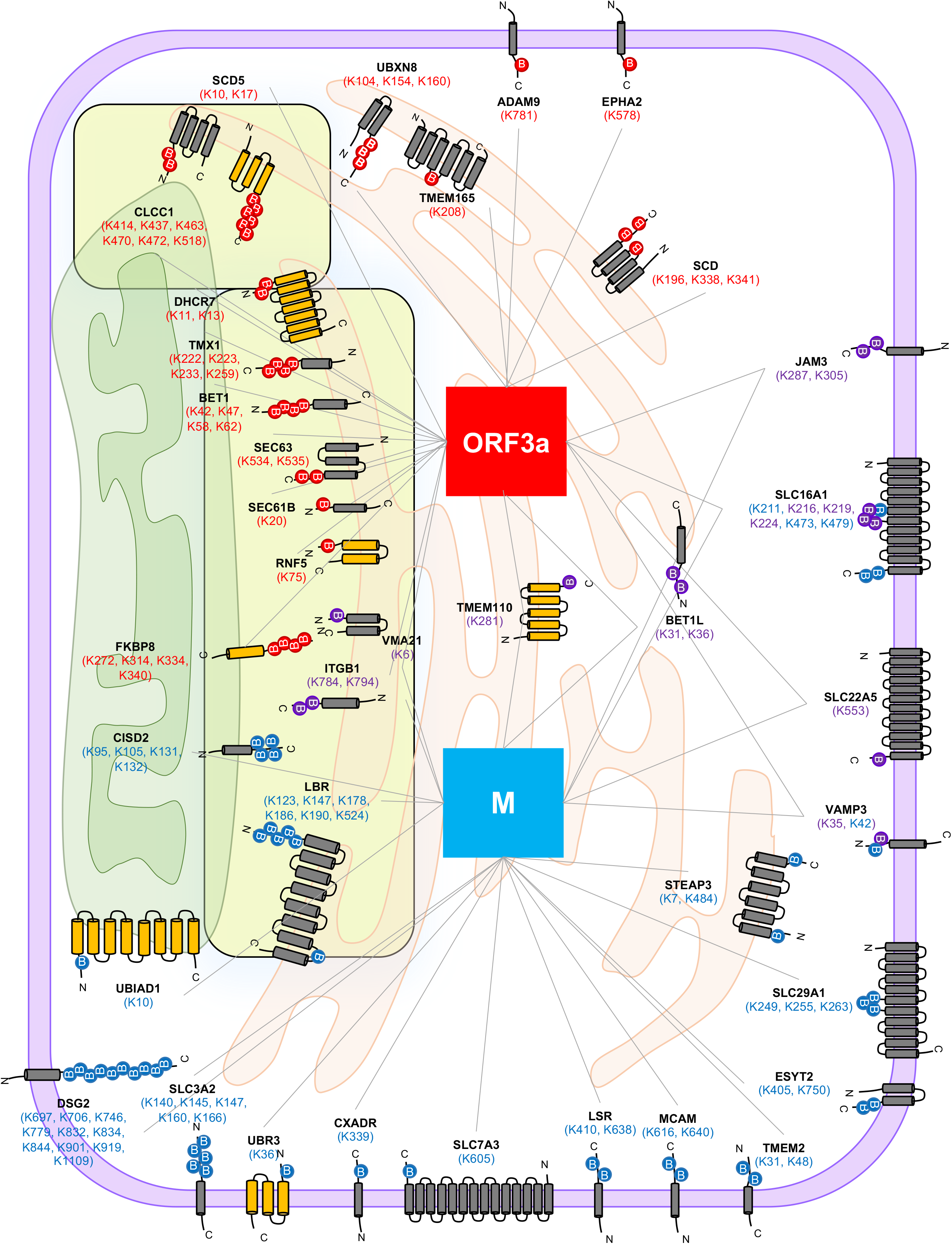
Proposed membrane topologies of the integral membrane proteins in ORF3a and M’s interactome list. Biotin-labeled sites by TurboID-GBP (cytosolic side) were colored according to their samples (red: observed in ORF3-GFP samples; blue: observed in M-GFP samples; purple: observed both in ORF3a and in M samples). Newly proposed membrane topologies are shown in yellow. Detailed information is given in **Figure S8** and **Dataset S1**.

It is noteworthy that this detailed topological information can be only obtained by direct detection of biotinylated peptides, however, conventional methods that overlook these biotin-PTM signatures cannot report it. We believe that our data provide detailed information on the domains heading toward cytosolic domain of ORF3a and M proteins at the membrane, which lays foundation for further studies on host membrane proteins. Notably, among the biotinylated membrane proteins, several lysosome-associated proteins (i.e., VPS16, VMA21, ITM2C, TMEM165, and RAB7A) were identified in the ORF3a interactome. This finding is consistent with recent reports that ORF3a hijacks the HOPS complex and RAB7, which are necessary for membrane contact between autophagosomes and lysosomes for autolysosome formation (Miao et al., 2021; Zhang et al., 2021). Our result implies that ORF3a may dynamically localize from the MAM to the autophagosome and autolysosome because autophagosome formation is also known to be initiated at the MAM (Hamasaki et al., 2013).

### RNF5 ubiquitinylate ORF3a of SARS-CoV-2

In regard to the ORF3a interaction with membrane proteins at the autophagosome and lysosome, RNF5, which is strongly biotinylated by *Spot-TurboID* (**Figure 6A**), interests us because it is an ER-anchored E3 ubiquitin-protein ligase involved in regulating the autophagic process during bacterial infections (Kuang et al., 2012). Interestingly, RNF5 is involved in the inflammatory response (Li et al., 2019), and it regulates the antiviral response by degrading an outer mitochondrial protein, MAVS (Zhong et al., 2009; Zhong et al., 2010), which implies that RNF5 might function at the MAM. Indeed, RNF5 was identified as an MAM-resident E3 ubiquitin ligase in our recent study (Kwak et al., 2020).

**Figure 6.**
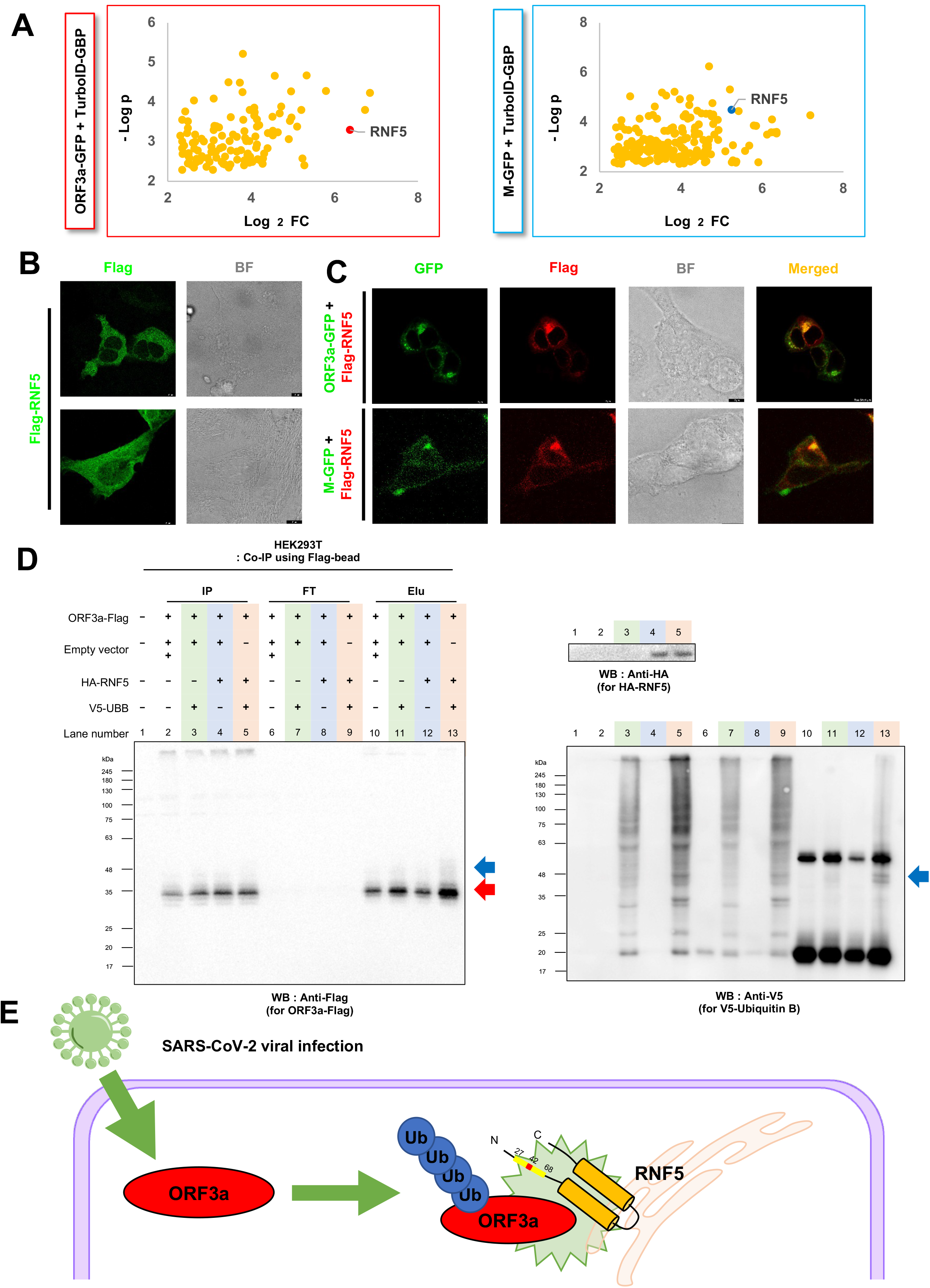
RNF5 ubiquitinylate ORF3a of SARS-CoV-2. **(a)** Highlighted RNF5 in ORF3a interactome (left) and in M interactome (right) volcano plots shown in **Figure 4b**. **(b)** Confocal microscopy images of Flag-RNF5 in HEK293-AD cells. Flag-RNF5 was visualized by anti-Flag antibody (AF488-conjugated, GFP fluorescence channel). Scale bars: 10 μm. **(c)** Confocal microscopy images of vPOI-GFP with Flag-RNF5. Flag-RNF5 was visualized by anti-Flag antibody (AF647-conjugated, Cy5 fluorescence channel). BF: bright field, Scale bars: 10 μm. (**d**) Western blot analysis of ORF3a ubiquitination by RNF5 in HEK293T cells. Anti-Flag and Anti-V5 were utilized for ORF3a-Flag and V5-UBB, respectively. Red arrow marks the unmodified molecular weight of ORF3a-Flag. Blue arrows mark ubiquitinylated ORF3a-Flag. Same molecular weight bands were detected in both anti-Flag and anti-V5 blot results. Additional blot results are shown in **Figure S9d-e** and **Figure S10**. **(e)** Proposed model of RNF5-mediated ubiquitination of ORF3a of SARS-CoV-2. Zinc finger domain of RNF5 (27–68 aa) is highlighted with yellow box, and 42 aa is an essential site for activity of E3 ubiquitin ligase.

Since the molecular function of RNF5 with SARS-CoV-2 is not fully characterized yet, we conducted further studies to reveal its functional relationship with SARS-CoV-2 proteins. From confocal microscopy experiments, we observed that cytoplasmic dispersed Flag-conjugated RNF5 was largely altered by ORF3a and M expression and co-localized with the respective viral proteins (**Figure 6B-C, S9A**). This result indicates that RNF5 might be activated by ORF3a and M protein expression. From western blot analysis after Co-IP, we observed that overexpression of RNF5 induced further ubiquitination of ORF3a protein (**Figure 6D, S9D-E**). It is noteworthy that overexpression of a mutant RNF5(C42S) whose ubiquitin-transferring ring finger domain is not functional owing to C42S mutation (Zhong et al., 2009) did not induce ubiquitination of ORF3a (**Figure S10**). These results indicate that RNF5 is involved in the ubiquitination of viral proteins and their interacting host proteins (**Figure 6E**).

In our study, results of western blot analysis revealed increased expression of ORF3a and M following MG132 treatment (**Figure S9B-C**). Thus, we propose that RNF5 is responsible for ubiquitination and degradation of ORF3a and M proteins derived from SARS-CoV-2. Our result is in a good agreement with a recent study that selected RNF5 as a specific ubiquitin ligase for viral proteins of SARS-CoV-2 from genome-wide screening (Zhen Yuan, 2021). Together with our result, we expect RNF5 to be an effective target for the development of anti-viral therapeutics for SARS-CoV-2.

### SARS-CoV-2 ORF3a influences ER–mitochondria contact sites and the secreted proteins of host cells

Interestingly, we also observed that many of the ORF3a and M interactome proteins overlapped with our recently identified MAM proteins. We have identified 115 local resident proteins using Contact-ID, which utilized the reconstituted biotinylating activity of split BioID fragments at the MAM, which is an ER and mitochondrial contact site (Kwak et al., 2020). Twenty-eight of the proteins that filtered out from the ORF3a interactome (23.9%) and the 24 from the M interactome (12.6%), overlapped with the 115 proteins previously identified to be a part of the MAM proteome. A total of 18 proteins overlapped between ORF3a, M, and the MAM proteome (**Figure 7A**). The complete list of proteins in the ORF3a and M proteomes that overlap with the MAM proteome, classified by function, are given in **Figures 7B** and **7C**, respectively. Since many of the ORF3a interactome proteins overlapped with those from the MAM proteome rather than the M interactome, we questioned whether ORF3a could more strongly hinder MAM formation.

**Figure 7.**
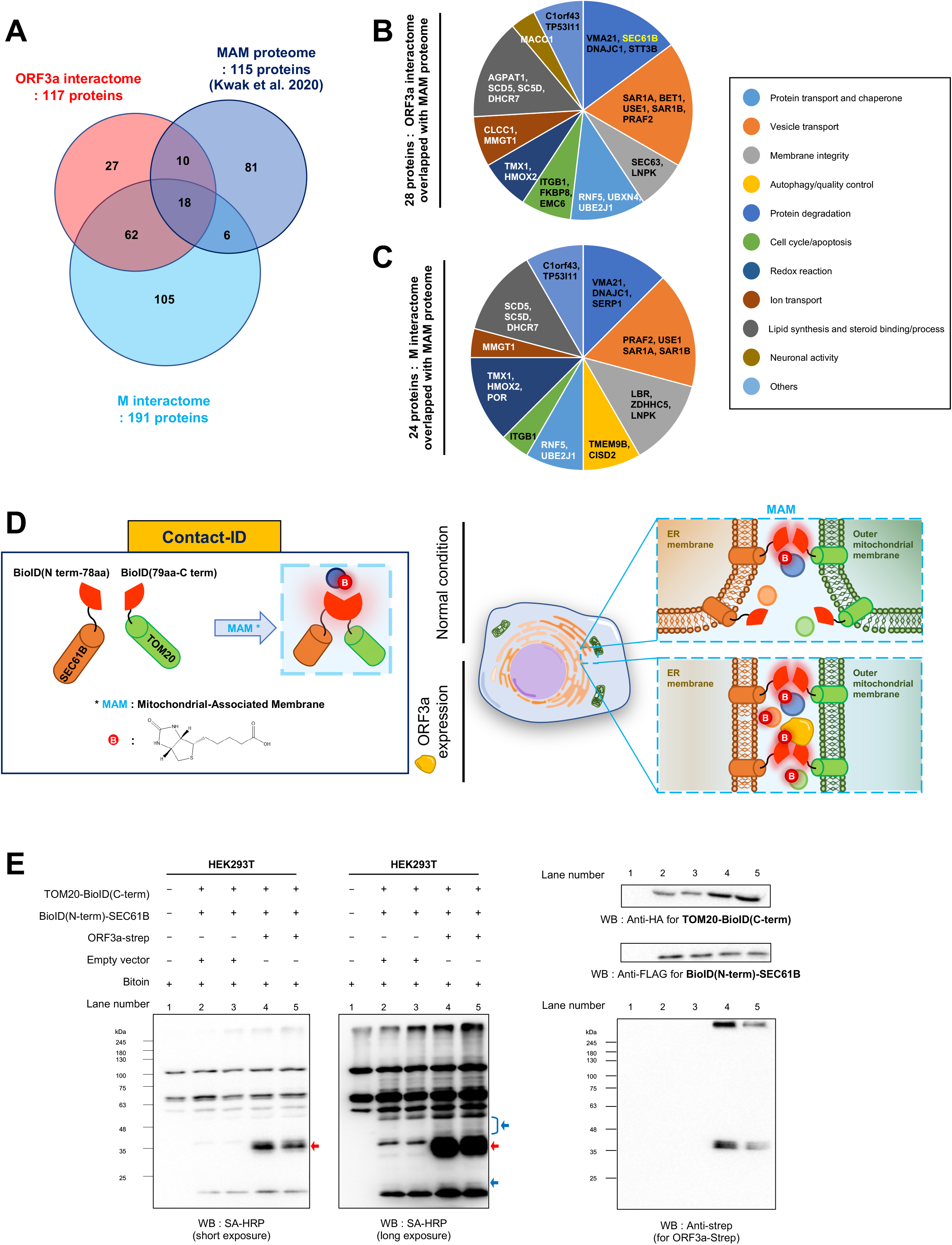
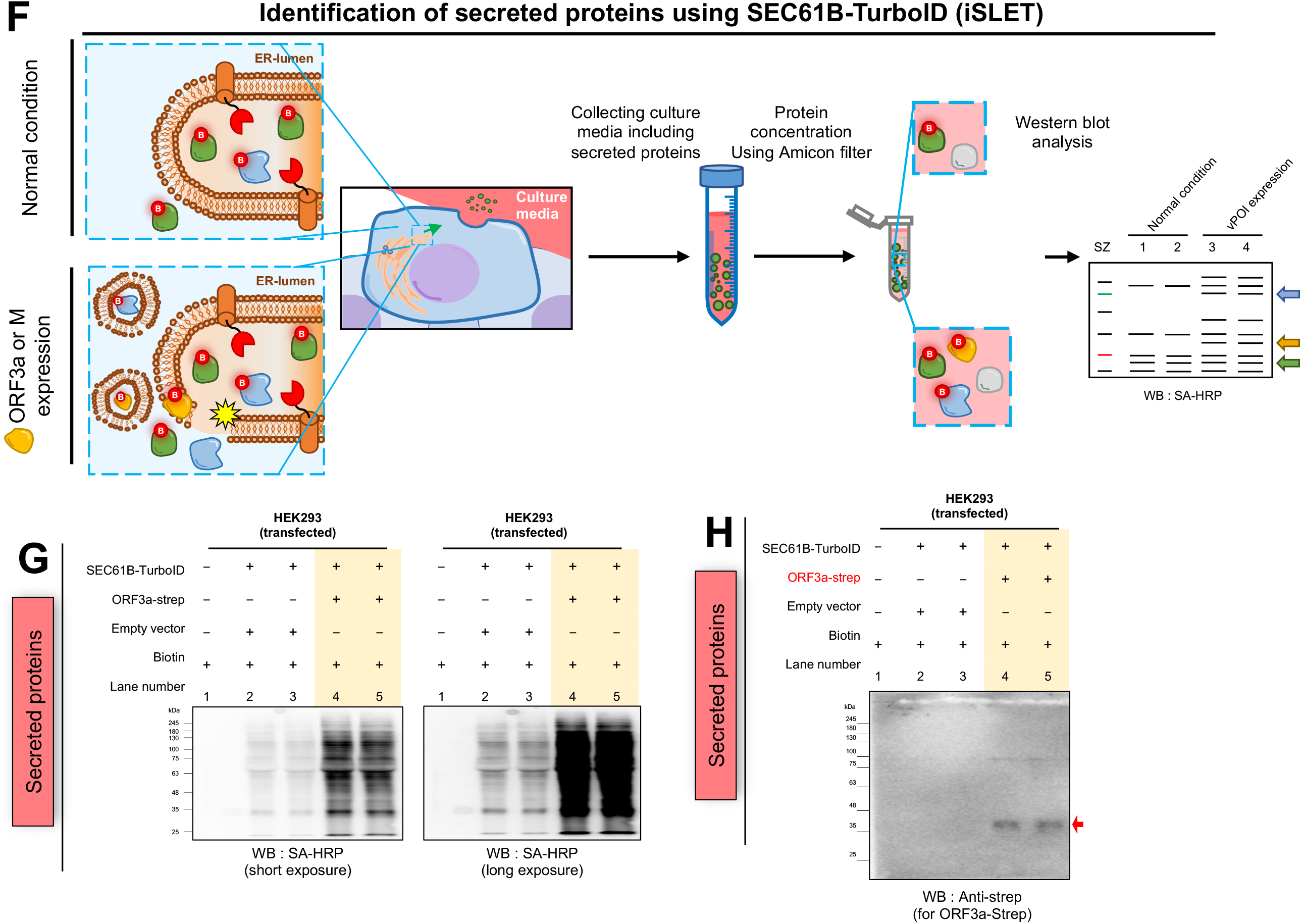
ORF3a remodeled local proteome at the MAM. **(a)** Local protein overlaps between interactomes of ORF3a (117 proteins), M (191 proteins), and MAM proteome (115 proteins). **(b)** Pie chart of functionally classified 28 overlapped proteins between ORF3a interactome and MAM proteome. **(c)** Pie chart of functionally classified 24 overlapped proteins between M interactome and MAM proteome. **(d)** Scheme of Contact-ID assay to check the MAM proteome changes by ORF3a expression. **(e)** SA-HRP western blotting results of Contact-ID with or without ORF3a co-expression. Biotinylated ORF3a (lane 4, 5) are marked with red arrow. Additional biotinylated proteins (lane 4, 5) are marked with blue arrows. Raw images are shown in **Figure S11. (f)** Scheme of *in situ* biotinylation of secreted protein (iSLET) by SEC61B-TurboID. **(g)** SA-HRP western blotting results of secreted biotinylated proteins from SEC61B-TurboID expressed HEK293 cells with or without co-expression of ORF3a. Raw images are shown in **Figure S13a. (h)** Anti-strep western blot results of the same samples as (**b).** Bands of ORF3a-strep (lane 4, 5) are marked with a red arrow. Raw images are shown in **Figure S13c.**

To address this, we used the Contact-ID tool to monitor changes in the proteomic composition at the MAM in response to ORF3a expression (**Figure 7D**). (Contact-ID generated more biotinylated proteins in the ORF3a-expressing sample compared to that in the control sample (blue arrows in **Figure 7E; Figure S11A-D**), which implies that ORF3a may lead to a considerable change of the proteomic landscape at the MAM. We also found that ORF3a was biotinylated by Contact-ID (red arrows in **Figure 7E**). Thus, we are currently conducting further investigations to analyze which proteins are recruited to the MAM in response to ORF3a expression. Similar to ORF3a, we observed that M was also biotinylated by Contact-ID (**Figure S12**).

We speculate that the expression of vPOI may change the secretory proteome because our EM results showed that ORF3a and M largely perturb the endomembrane system and many viral infections have been known to affect secretory proteome profiles (Lietzén et al., 2019). To investigate alterations in the host cell secretome, we used recently developed method dubbed iSLET (*in situ* secretory protein labeling via ER-anchored TurboID) (Kim et al., 2020). In the iSLET method, SEC61B-TurboID is utilized to biotinylate secreted proteins at the ER luminal side near to the SEC61 translocon channel since most secreted proteins transit through the ER lumen via the SEC61B translocon channel. Secretory proteins biotinylated in SEC61B-TurboID-expressing cells can be detected in the culture media or mouse bloodstream. In this study, we utilized the iSLET method to assess whether the secretome profiles of the host cells changed in response to ORF3a and M expression (**Figure 7F**).

In the SA-HRP western blot results, the amounts of biotinylated proteins in the culture media of cells co-expressing SEC61B-TurboID and ORF3a were substantially higher compared to that of the control sample lacking ORF3a expression (**Figure 7G, S13A-B**). In addition, we found ORF3a in the culture media of the cells (red arrows in **Figure 7H, S13C**). In whole cell lysates, ORF3a was also biotinylated by SEC61B-TurboID (**Figure S14**). Similar to ORF3a, we observed that the amounts of biotinylated proteins increased in the secretome of M-expressed cells (**Figure S15A**), and M protein was also secreted from cells (**Figure S15B**) and was biotinylated by SEC61B-TurboID (**Figure S15C**).

These results imply that ORF3a and M may upregulate cargo secretion from the ER and there may be a specific sorting pathway for the secretion of ORF3a in the ER lumen, as has been shown for other secreted viral proteins (Alefantis et al., 2005). In the ORF3a interactome, SAR1A/B and USE1 are known to play essential roles in cargo secretion at the ER-Golgi interface (Petrosyan et al., 2015) and further investigation is required to identify whether the interaction of these vesicle transport proteins with ORF3a could induce ER-derived vesicle formation and changes in the secretome profile.

## Discussion

In this study, we demonstrated that ORF3a and M perturb the integrity of endomembrane systems in host cells, thereby resulting in devastating effects. As shown in other viruses (Dengue, porcine epidemic diarrhea, and Zika viruses) that utilize the ER for replication (Lee et al., 2018; Monel et al., 2017; Wang et al., 2012a; Zou et al., 2019), including SARS-CoV (Fung et al., 2014), our data show that ORF3a and M of SARS-CoV-2 largely affect membrane integrity and the proteomic landscape at the ER. Although we employed ectopic expression method of ORF3a and M protein constructs to ensure biosafety, a recent study shows that ectopic expression of ORF3a of SARS-CoV-2 is enough to show pathogenesis of SARS-CoV-2 in *Drosophila* (Shuo Yang, 2020). Since some of the proteins we identified, including RNF5, are also highlighted as viral host proteins in other studies of SARS-CoV-2 (Zhen Yuan, 2021), we believe that our interactome data reflects physiological networks of the ORF3a and M proteins of SARS-CoV-2.

During our interactome analyses of ORF3a and M, we observed a notable overlap with the MAM proteome (Kwak et al., 2020). Among the overlaps, VAP-A and VAP-B interact with NS5A and NS5B to promote viral replication and protein assembly of hepatitis C virus at the MAM (Hamamoto et al., 2005). We confirmed that RNF5 (Kuang et al., 2012), co-localized with ORF3a and induced ubiquitination of ORF3a (**Figure 6**). This result is consistent with previous studies that showed ubiquitination of SARS-CoV-2 ORF3a (Cao et al., 2021) and RNF5-induced ubiquitination of SARS-CoV-2 M (Zhen Yuan, 2021). Whether overexpression or chemical activation of RNF5 could be used as a novel anti-viral or anti-inflammatory strategy for SARS-CoV-2 warrants further investigations.

In our study, RNF5 was identified as part of the ORF3a interactome, whereas previously reported interactome mapping approaches such as conventional proximity labeling technique (Samavarchi-Tehrani et al., 2020) and affinity-purification method (Gordon et al., 2020) could not identify RNF5 in the ORF3a interactome although their selections contain comparable or more numbers of findings in the list. Thus, our *Spot-TurboID* method is superior to previous methods for identifying proteins of interest in an interactome.

Since MAM is a cholesterol-enriched intracellular lipid raft (Area-Gomez et al., 2012), several proteins associated with lipid metabolism and transport are observed at this domain (Kwak et al., 2020). Notably, among 119 proteins identified in ORF3a interactome in this study, 34 proteins were earlier characterized as lipid binding proteins (LBP) (Niphakis et al., 2015) and 15 proteins were reported at the MAM (Kwak et al., 2020) (**Dataset S1**). Thus, we speculate that ORF3a forms neo-organelles (i.e. CM and DMV) by altering ultrastructure and lipid contents of MAM since dynamic lipid transport at the MAM (Phillips and Voeltz, 2016) can supply lipid molecules for the formation of autophagosomal membrane formations (Hamasaki et al., 2013). Since several lysosome-associated membrane proteins (e.g. VPS16, RAB7A) are observed in our ORF3a interactome as also observed in other ORF3a interactome list using affinity purification method (Stukalov et al., 2021), consistent with recent reports that ORF3a is localized at the late endosome and lysosome (Miao et al., 2021; Zhang et al., 2021), we postulated that these ORF3a-associated vesicles (e.g., CM and DMV) might be interconnect or fused with the lysosome at some points.

Additionally, MAM is a site of active calcium transport from the ER to the mitochondria; a recent study characterized ORF3a of SARS-CoV-2 as Ca^2+^ and K^+^ channels. Thus, we postulate that ORF3a may derange the calcium transport, which in turn, can negatively impact MAM function. Since abnormal MAM formations correlate with various metabolic diseases such as type 2 diabetes (Tubbs et al., 2018) and neurodegenerative diseases, including Alzheimer’s disease, Parkinson’s disease, amyotrophic lateral sclerosis (Schon and Przedborski, 2011), and Wolfram syndrome (Delprat et al., 2018), further studies might be required to monitor the possible progression of these MAM-related metabolic diseases in SARS-CoV-2-infected patients (Rahman et al., 2020).

In addition, we also found that many of the proteins in the ORF3a and M interactomes are targeted by approved drug molecules, which may be useful for drug repurposing therapeutic strategies. The druggable proteins targeted by previously approved drug molecules are listed in **Supplementary Table 2-3**.

In this study, we also provide evidence that our *Spot-TurboID* workflow using the GFP/GBP binding system (vPOI-GFP/TurboID-GBP) is effective for interactome mapping of viral proteins in live cells. Since several viral proteins are initially cloned with GFP for the imaging experiments typically performed in the initial stages of many virus studies, we expect that our TurboID-GBP could be an effective “plug-and-playable” component for rapid host interactome analysis with various GFP-tagged viral proteins, without cloning additional constructs for PL experiments. This is especially relevant in situations where time is a constraint, such as that in the current COVID-19 pandemic.

## Supporting information

Data S1. ORF3a and M interactive

## Resource Availability

### Lead Contact

Further information and request of reagents or resources should be directed Contact, Dr. Rhee H. W.

### Materials Availability

Upon reasonable requests, unique reagents utilized in this paper could be provided.

### Data and Code Availability

The raw LC-MS data generated from this study have been deposited on ProteomeXchange Consortium (accession no. PDX022355, User name, Password: jvuXMLgS).

## Materials

| REAGENT or RESOURCE | SOURCE | IDENTIFIER |
| --- | --- | --- |
| Antibodies |  |  |
| V5 Tag Monoclonal Antibody (mouse) | Invitrogen | #R960-25 |
| GFP Monoclonal Antibody (mouse) | Invitrogen | #MA5-15256 |
| Anti-StrepMAB-Classic, purified, lyophilized, (mouse) | IBA | #2-1597-001 |
| HA Tag Polyclonal Antibody | Thermofisher | #71-5500 |
| FLAG rabbit Polyclonal Antibody | Sigma aldrich | #F7425 |
| Ubiquitin Antibody | Santa Cruz biotechnology | #SC-166553 |
| Alexa Fluor 568 IgG mouse | Molecular probes | #A11004 |
| Streptavidin, Alexa Fluor 647 conjugate | Invitrogen | #S21374 |
| Goat Anti-Mouse IgG (H + L)-HRP conjugate | Bio-rad | #1706516 |
| Anti-rabbit IgG, HRP-linked Antibody | Cell signaling | #7074S |
| Streptavidin-HRP | Thermofisher | #21126 |
| Anti-Flag affinity gel beads | Bimaker | #B23102 |
| 4% paraformaldehyde | CHEMBIO | #CBPF-9004 |
| HyClone Dulbecco's Modified Eagle Medium (DMEM) with high glucose: Liquid | Cytiva | #SH30022.01 |
| FBS | Atlas Biologicals | #EF-0500-A |
| biotin | Alfa aesar | #A14207 |
| doxycycline | Sigma aldrich |  |
| RIPA lysis buffer | ELPISBIO | #EBA-1149 |
| Protease inhibitor cocktail | Invitrogen | #78438 |
| Reagent for mass preparation |  |  |
| Sodium azide | Alfa aesar | #14314 |
| Sodium ascorbate | Sigma aldrich | #A4034 |
| Trolox | Sigma aldrich | #238813 |
| Urea | Sigma aldrich | #U5378 |
| 20X TBS | Thermofisher | #28358 |
| Acetone | Sigma aldrich | #650501 |
| Ammonium bicarbonate | Sigma aldrich | #A6141 |
| Sodium dodecyl sulfate | Sigma aldrich | #436143 |
| TPCK-Trypsin | Thermofisher | #20233 |
| Dithiothreitol | Sigma aldrich | #43819 |
| Iodoacetoamide | Sigma aldrich | #I1149 |
| Trifluoroacetic acid | Sigma aldrich | #T6508-10AMP |
| Formic acid | Thermofisher | #28905 |
| Streptavidin-coated magnetic beads | Pierce | #88817 |
| Equipment |  |  |
| Speed-vac |  |  |
| Thermomixer |  |  |

## Methods

### Cell lines

HEK293 cells and HEK293T cells were gift from Professor Kim, H. at Ulsan National Institute of Science and Technology (Korea). SEC61B-V5-TurboID stably expressed HEK293 Flip-in T-rex cells were from the past paper. HeLa cells and A549 cells were from Laboratory of Dr. Mun at Korea Brain Research Institute (Korea). Cell lines were cultured according to the manufacturer’s instructions.

### Expression Plasmids

Genes with epitope tag (i.e. V5, FLAG, HA, twin strep) were cloned into pCDNA3, pCDNA3.1, and pCDNA5 using digestion with enzymatic restriction sites and ligation with T4 DNA ligase. After digesting PCR products and cut vectors simultaneously, they were ligated. CMV promoter made genes expressed possible in mammalian cells. Table S1 summarized constructs information. Templates of SARS-CoV-2 were obtained from Professor Kim, Ho Min (KAIST) and Professor Kim, V. Narry (SNU).

### Cell Culture and Transfections

Cell lines were examined for morphology and confluence using a microscope. Cells were incubated with in Dulbecco’s Modified Eagle Medium (Cytiva, cat. No., SH30022.01) supplemented with 10% fetal bovine serum (Atlas biologicals, cat. No., EF-0500-A) in a 37 °C incubator 5% CO_2_ (v/v). Genes were introduced into cells using PEI, when 60–80% cell confluence.

### Stable Expression of SEC61B-V5-TurboID Cell Line (Flip-in HEK293T-Rex)

SEC61B-V5-TurboID expression was induced by treatment of 5 ng/mL doxycycline (Sigma Aldrich). For transient expression of other constructs, transfection using PEI could do the same time with treatment of doxycycline.

### Biotin Labeling in Live Cells

Using transient transfection reagent, PEI, genes including *TurboID* or *BioID* were introduced into HEK293-AD, HEK293 or HEK293T cells. After 16–24 h, the medium was changed to 500 μL (for 24 well plate) or 1 mL (for 12 well plate) of fresh growth medium containing 50 μM biotin (Alfa aesar, cat. No. A14207). These cells were incubated at 37 °C and 5% CO_2_ for 30 min for using TurboID (or 16 h for using split BioID) in accordance with the previously published protocols. Next, the reaction was stopped by washing, two times, with Dulbecco’s phosphate-buffered saline (DPBS). After, lysis was followed for western blot analysis or mass sampling, or fixation was followed for fluorescence microscope imaging.

Using transient transfection reagent, PEI, genes including APEX2 were introduced into HEK293-AD, HEK293 or HEK293T cells. After 16–24 h, the medium was changed to 500 μL (for 24 well plate) or 1 mL (for 12 well plate) of fresh growth medium containing 250 μM desthiobiotin-phenol. These cells were incubated at 37 °C and 5% CO_2_ for 30 min in accordance with the previously published protocols (Lee et al., 2017). H2O2 (1 mM; Sigma Aldrich, cat. No., STBJ2658) was added to each well for 1 min at room temperature. Next, this reaction was terminated by washing, two times, with Dulbecco’s phosphate-buffered saline (DPBS) containing 5 mM Trolox (Sigma Aldrich, cat. No., 238813-25G), 10 mM sodium azide (Alfa Aesar, cat. No., N07E043), and 10 mM sodium ascorbate (Sigma Aldrich, cat. No., A4034-100G). After, lysis was followed for western blot analysis or mass sampling, or fixation was followed for fluorescence microscope imaging.

### Fluorescence Microscope Imaging

Cells were split on the coverslips (thickness no. 1.5 and radium: 18 mm) for microscope imaging.vPOI-GFP expressed cells were fixed using 4% paraformaldehyde solution (CHEMBIO, cat. No. CBPF-9004) at room temperature for 10–15 min. Cells were then washed with Dulbecco’s phosphate-buffered saline (DPBS) two times. To detect vPOI-GFP expression, washed cells were maintained in DPBS on 4 °C refrigerator for imaging by Leica (NICEM in Seoul National University, Korea) with objective lens (HC PL APO 100x/1.40 OIL), White Light Laser (WLL, 470–670 nm, 1 nm tunable laser), HyD detector and controlled by LAS X software.

Co-expressed or biotin-labeled cells were fixed with 4% paraformaldehyde solution (CHEMBIO, cat. No. CBPF-9004) in DPBS at room temperature for 10–15 min. Cells were then washed with DPBS, two times, and permeabilized with cold methanol at −20 °C for 10 min. Cells were washed again, two times, with DPBS and blocked for 30 min with 2% BSA in DPBS at room temperature (25°C). To detect expression pattern or biotinylation pattern, the primary antibody, such as anti-V5 (Invitrogen, cat. No. R960-25, 1:5,000 dilution), was incubated for 1 h at room temperature. After washing two times with TBST, cells were simultaneously incubated with secondary Alexa Fluor 568 IgG mouse (Molecular probes, cat. No. A11004, 1:1,000 dilution) and Streptavidin, Alexa Fluor 647 conjugate (Invitrogen, cat. No. S21374, 1:1,000 dilution) for 30 min at room temperature. Cells were then washed two times with TBST and maintained in DPBS at 4 °C for imaging using Leica (NICEM in Seoul National University, Korea). Pearson correlation was conducted using image J software program (NIH).

### Co-immunoprecipitation

Constructs were transfected using PEI. After more than 16 h, samples were lysed using RIPA buffer (ELPISBIO, cat. No. EBA-1149) including 1× protease inhibitor cocktail (Invitrogen, cat. No. 78438) for 10 min at room temperature. After transferring these lysates to e-tube, samples were vortexed for 2–3 min at room temperature. After centrifugation at 15,000 × *g* for 10 min at 4 °C, supernatant was collected. These samples were incubated with already washed Anti-Flag affinity gel beads (Bimaker, cat. No.B23102) for 2 h at 4 °C. Then they were washed more than three times and eluted using 1× SDS-PAGE loading buffer at 95 °C for 5 min.

### Obtaining Secreted Proteins in Cell Culture Media

After biotin labeling of HEK293 cells expressing SEC61B-TurboID, cells were incubated with no FBS DMEM for 16 h. After collecting these cell culture media in 15 mL tube, these samples were concentrated using amicon filter (Millipore, cat. No. UFC501096) at 12,000 × *g* at 4 °C for 20 min. Concentrated proteins were collected using 8 M urea (Sigma Aldrich, cat. No. U5378). These samples were used for immunoblotting.

### Immunoblotting

Transfected and biotinylated cell samples were lysed using RIPA buffer (ELPISBIO, cat. No. EBA-1149) including 1× protease inhibitor cocktail (Invitrogen, cat. No. 78438) for 10 min at room temperature. After transferring these lysates to e-tube, samples were vortexed for 2–3 min at room temperature. Using centrifugation at 15,000 × *g* for 10 min at 4 °C, samples were clarified. They were boiled with 1× SDS-PAGE loading buffer at 95 °C for 5 min. After resolving protein samples using SDS-PAGE and transferring protein samples to membrane, immunoblotting analysis was performed using antibodies. Membranes were blocked for 30 min with 2% skim milk solution at room temperature. After washing three times with TBST for 5 min each, at room temperature, primary antibodies, namely anti-V5 (Invitrogen, cat. No. R960-25, 1:10,000 dilution), anti-GFP (Invitrogen, cat. No. MA5-15256, 1:3,000 dilution), anti-HA (Thermofisher, cat. No. 71-5500, 1:10,000 dilution), anti-FLAG (Sigma Aldrich, cat. No. F7425, 1:10,000 dilution), and anti-strep (IBA, cat. No. 2-1597-001, 1:1,000 dilution), was incubated for 16 h at 4 °C. After washing three times with TBST for 5 min at room temperature, secondary antibodies, namely mouse-HRP (Bio-rad, cat. No. 1706516, 1:3,000 dilution) and rabbit-HRP (Cell signaling, cat. No. 7074S, 1:3,000 dilution), or SA-HRP (Thermofisher, cat. No. 21126, 1:10,000 dilution) were incubated for 1 h at room temperature. After washing three times with TBST for 5 min at room temperature, results of immunoblotting assay were obtained using ECL solution using GENESYS program. Line scan analysis was performed using image J software program (NIH). After subtraction of intensity value at background, protein bands from top to bottom of each lane were putted as x-axis and intensity of bands from top to bottom of each lane was putted as y-axis.

### EM imaging and CLEM

To observe the DAB-stained cells, cells were cultured in 35-mm glass grid-bottomed culture dishes (MatTek life sciences, MA, USA) to 30–40% confluency. Then cells were transfected with DNA plasmids using Lipofectamin 2000 (Life Technologies, Carlsbad, CA, USA). Next day, cells were fixed with 1% glutaraldehyde (Electron Microscopy Sciences, cat. No. 16200) and 1% paraformaldehyde (Electron Microscopy Sciences, cat. No. 19210) in 0.1 M cacodylate solution (pH 7.0) for 1 h at 4 °C. After washing, 20 mM glycine solution was used for quench unreacted aldehyde. DAB staining (DAB, Sigma, cat. No. D8001) took approximately 20−40 min until a light brown stain was visible under an inverted light microscope. DAB stained cells were post-fixed with 2% osmium tetroxide in distilled water for 30 min at 4 °C and *en bloc* in 1% uranyl acetate (EMS, USA, cat. No. 22400) overnight and dehydrated with a graded ethanol series. The samples were then embedded with an EMBed-812 embedding kit (Electron Microscopy Sciences, USA, cat. No. 14120) and polymerized in oven at 60 °C. The polymerized samples were sectioned (60 nm) with an ultramicrotome (UC7; Leica Microsystems, Germany), and the sections were mounted on copper slot grids with a specimen support film. Sections were stained with uranyless (Electron Microscopy Sciences, cat. No. 22409) and lead citrate (Electron Microscopy Sciences, cat. No. 22410) then viewed on a Tecnai G2 transmission electron microscope (ThermoFisher, USA).

For the CLEM observation, the ORF3a-GFP, M-mCherry and M-GFP transfected cells were imaged under a confocal light microscope (Ti-RCP, Nikon, Japan) in living cell, and cells were fixed with 1% glutaraldehyde and 1% paraformaldehyde in 0.1 M cacodylate solution (pH 7.0). After washing, cells were dehydrated with a graded ethanol series and infiltrated with an embedding medium. Following embedment, 60 nm sections were cut horizontally to the plane of the block (UC7; Leica Microsystems, Germany) and were mounted on copper slot grids with a specimen support film. Sections were stained with uranyl acetate and lead citrate. The cells were observed at 120 kV in a Tecnai G2 microscope (ThermoFisher, USA). Confocal micrographs were produced as high-quality large images using PhotoZoom Pro 8 software (Benvista Ltd., Houston, TX, USA). Enlarged fluorescence images were fitted to the electron micrographs using the Image J BigWarp program.

### Proteome Digestion & Enrichment of Biotinylated Peptides

For mass sampling, HEK293T cells were split in three T75 flasks for triplicate samples per each condition. For transiently co-expressed two constructs, vPOI-GFP and TurboID-GBP, cells were grown at 70–80% confluence and transfected with wanted constructs using PEI. After 16 h (transfection), 50 μM biotin (Alfa aesar, cat. No. A14207) were treated for 30 min in 37 °C incubator. After biotin labeling, these cells were washed 3–4 times with cold DPBS and lysed with 1.5 mL 2% SDS in 1× TBS (25 mM Tris, 0.15 M NaCl, pH 7.2, Thermoscientific, cat. No. 28358) containing 1× protease inhibitor cocktail. These lysates were underwent ultrasonication (Bioruptor, diagenode) 3–4 times for 15 min in a cold room. For removing free biotin, 5–6 times the sample volume of cold acetone stored at −20 °C (Sigma Aldrich, cat. No. 650501) was mixed with lysates and stored at −20 °C for least 2 h. These samples were centrifuged at 13,000 × *g* for 10 min at 4 °C. Supernatant solution was gently discarded. Cold acetone, 6.3 mL, and 700 μL of 1× TBS were mixed with the pellet. These mixtures were then vigorously vortexed and stored at −20 °C for least 2 h or overnight. The samples were then centrifuged at 13,000 × *g* for an additional 10 min at 4 °C. Supernatant solution was gently discarded. After pellet was naturally dried for 3–5 min, it was resolubilized with 1 mL of 8 M urea (Sigma Aldrich, cat. No. U5378) in 50 mM ammonium bicarbonate (ABC, Sigma Aldrich, cat. No. A16141). Protein concentration was calculated using the BCA assay. Samples were vortexed at 650 rpm for 1 h at 37 °C using Thermomixer (Eppendorf) for denaturation. Samples were reduced in 10 mM dithiothreitol (Sigma Aldrich, cat. No. 43816) at 650 rpm for 1 h at 37 °C using Thermomixer (Eppendorf), while samples alkylation was accomplished in 40 mM iodoacetoamide (Sigma Aldrich, cat. No. I1149) at 650 rpm for 1 h at 37 °C using Thermomixer (Eppendorf). These samples were diluted 8 times with 50 mM ABC (Sigma Aldrich, cat. No. A16141). CaCl2 (Alfa aesar, cat. No. 12312) was added for the final concentration of 1 mM. Samples were digested using trypsin (Thermoscientific, cat. No. 20233) (50:1 w/w) at 650 rpm for 6–18 h at 37 °C using Thermomixer (Eppendorf). Insoluble materials were removed using centrifugation for 3 min at 10,000 × *g*. Streptavidin beads (300 μL; Pierce, cat. No. 88817) were washed using 2 M urea in 1× TBS for 3–4 times, were then added to samples, and then rotated for 1 h at room temperature. The flow-through fraction was not discarded, and beads were washed 2–3 times using 2 M urea in 50 mM ABC. After removing the supernatants, beads were washed using pure water and transferred to new tubes. After adding 500 μL 80% acetonitrile (Sigma Aldrich, cat. No. 900667), 0.2% TFA (Sigma Aldrich, cat. No. T6508) and 0.1% formic acid (FA, Thermoscientific, cat. No. 28905), heat such biotinylated peptides at 60 °C and mixed at 650 rpm. Supernatant without streptavidin beads was transferred to new tubes. This elution step was repeated at least 4 times. These total elution fractions were dried for 5 h using Speed-vac (Eppendorf). Before samples were injected to mass spectrometry, they were stored at −20 °C.

### Calculation of Similarity Value

Similarity value of each mass samples was calculated as previously reported. The correlation index, C_ij_, was introduced for calculation of similarity value. The Equation below shows how similarity value was calculated.

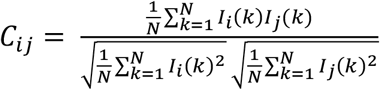

As the result of C_ij_ covers from 0 to 1, bigger value indicates more similarity, while smaller value indicates less similarity.

### LC-MS/MS Analysis of Enriched Peptide Samples

Analytical capillary columns (100 cm × 75 μm i.d.) and trap columns were packed in-house with 3 m Jupiter C18 particles (Phenomenex, Torrance). The long analytical column was placed in a column heater (Analytical Sales and Services) regulated to a temperature of 45 °C. NanoAcquity UPLC system (Waters, Milford) was operated at a flow rate of 300 nL/min over 2 h with linear gradient ranging from 95% solvent A (H_2_O with 0.1% formic acid) to 40% of solvent B (acetonitrile with 0.1% formic acid). The enriched samples were analyzed using Orbitrap Fusion Lumos mass spectrometer (Thermo Scientific) equipped with an in-house customized nanoelectrospray ion source. Precursor ions were acquired (m/z 300–1,500) at 120 K resolving power and the isolation of precursor for MS/MS analysis was conducted with a 1.4 Th. Higher-energy collisional dissociation (HCD) with 30% collision energy was used for sequencing with auto gain control (AGC) target of 1E5. Resolving power for acquired MS2 spectra was set to 30 k at with 200 ms maximum injection time.

### Processing MS data and Identification of Proteins

All MS/MS data were searched using MaxQuant (version 1.5.3.30) with Andromeda search engine at 10 ppm precursor ion mass tolerance against the SwissProt *Homo sapiens* proteome database (20,199 entries, UniProt (http://www.uniprot.org/)). The label free quantification (LFQ) and Match Between Runs were used with the following search parameters: Semi-trypic digestion, fixed carbaminomethylation on cysteine, dynamic oxidation of methionine, protein N-terminal acetylation with biotin labels of lysine residue. Less than 1% of false discovery rate (FDR) was obtained for unique labeled peptide and as well as unique labeled protein. LFQ intensity values were log-transformed for further analysis and missing values were filled by imputed values representing a normal distribution around the detection limit. The imputation of protein intensity were conducted using Perseus software program.

## ACKNOWLEDGMENTS

This work was supported by the National Research Foundation of Korea (NRF-2020K1A3A1A47110634, NRF-2019R1A2C3008463), the Organelle Network Research Center (NRF-2017R1A5A1015366) and a grant of the Korea Health Industry Development Institute(KHIDI) funded by the Ministry of Health & Welfare and Ministry of science and ICT, Republic of Korea (grant number : HU20C0326). H.W.R. was supported by Creative-Pioneering Researchers Program through Seoul National University. J.Y.M. and M.J. are supported by KBRI basic research program through Korea Brain Research Institute funded by Ministry of Science and ICT (Information & Communication Technology) (20-BR-01-09). EM data were acquired at Brain Research Core Facilities in KBRI. J.-S.K. are supported by IBS-R008-D1 of the Institute for Basic Science from the Ministry of Science and ICT of Korea.

## AUTHOR CONTRIBUTIONS

Y.B.L., J.Y.M., J.S.K., and H.W.R. conceived the project. M.J. and J.Y.M. contributed to EM imaging. J.K. and J.S.K. performed LC-MS/MS experiment and mass data processing. M.G.K. contributed to confocal imaging. C.K. contributed to preparation of mass samples. Y.B.L., J.Y.M., J.S.K., and H.W.R. wrote and edited the manuscript.

## SUPPLEMENTAL INFORMATION

Table S1-3.

Figures S1-S15.

Data S1. Result from mass analysis of ORF3a and M interactome

