## Supplementary information for "Endomembrane systems are reorganized by ORF3a and Membrane (M) of SARS-CoV-2"

**Table of Contents**

**Supplementary Figure 1-15------------------------------------------------------------------------------------S2-24**

**Supplementary Table 1-3-------------------------------------------------------------------------------------S25-29**

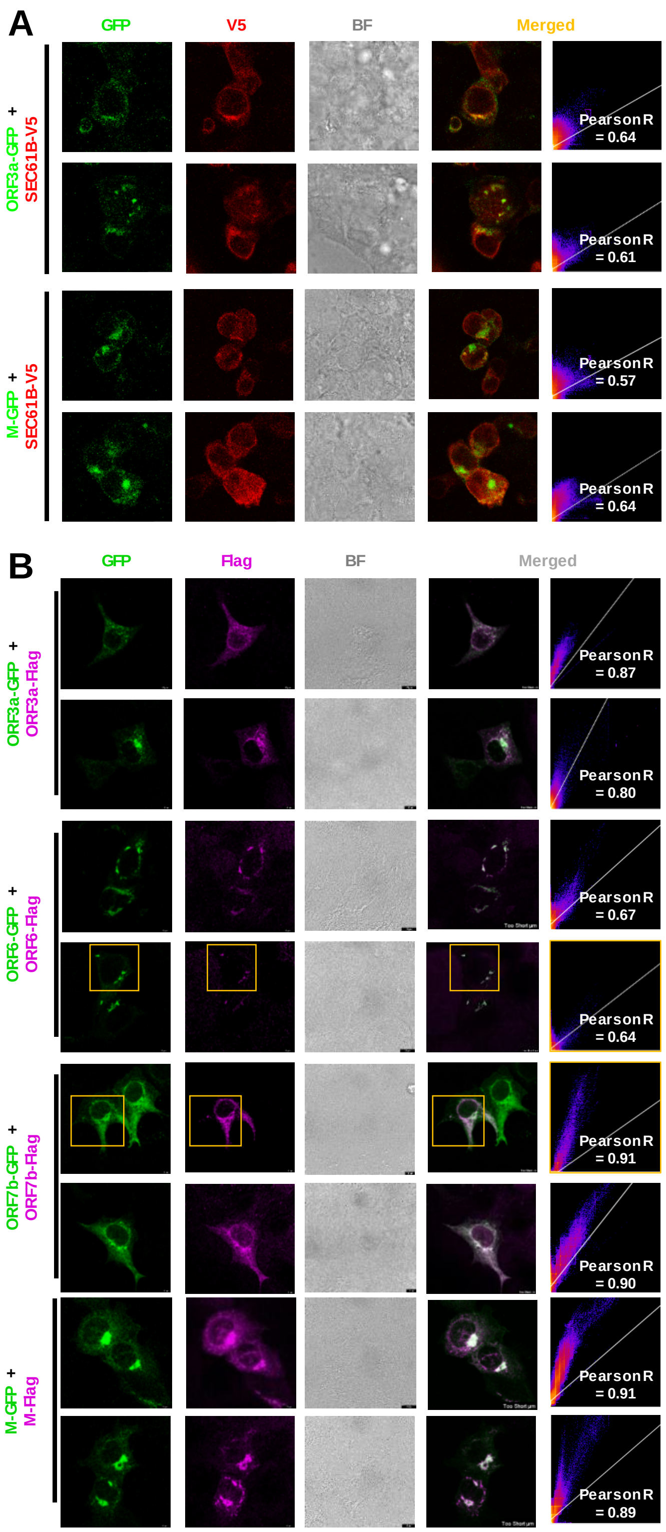


**Supplementary Figure 1. Additional confocal microscopy images of GFP-tagged vPOIs of SARS-CoV-2 (a)** Confocal microscopy images of ORF3a-GFP (left) and M-GFP (right) with SEC61B-V5-TurboID. SEC61B-V5-TurboID was visualized by Anti-V5 antibody (AF647-conjugated secondary antibody, Cy5 fluorescence channel). Pearson correlation values were calculated between different fluorescent channel images. BF: bright field, Scale bars: 10 μm. **(b)** Confocal microscopy images of co-expressed vPOI-GFP and vPOI-Flag. Flag-conjugated vPOIs (ORF3a, ORF6, ORF7b, M of SARS-CoV-2) were visualized by using Anti-Flag antibody (AF647-conjugated secondary antibody, Cy5 fluorescence channel) and GFP-conjugated vPOIs fluorescence was detected in GFP fluorescence channel. Pearson correlation values were calculated between different fluorescent channel images. BF: bright field, Scale bars: 10 μm.

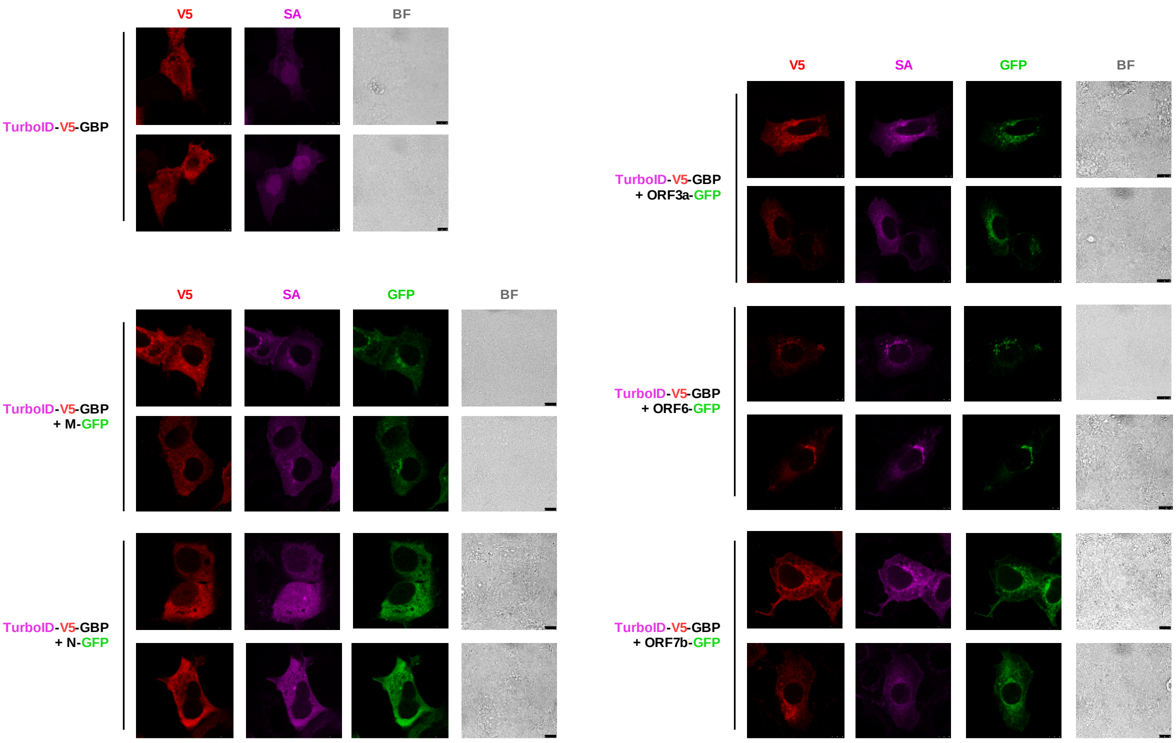


**Supplementary Figure 2. Additional Confocal images of Figure 2B.** Confocal microscopy images of GFP-conjugated vPOIs (M, N, ORF3a, ORF6 and ORF7b of SARS-CoV-2) and co-expressed TurboID-V5-GBP in HEK293-AD cells. TurboID-V5-GBP (TurboID-GBP) was visualized by anti-V5 antibody (AF568-conjugated, RFP fluorescence channel). and GFP fluorescence was detected in GFP fluorescence channel. Biotinylated proteins were visualized by AF647-conjugated streptavidin (Cy5 fluorescence channel). BF: bright field, Scale bars: 10 µm.

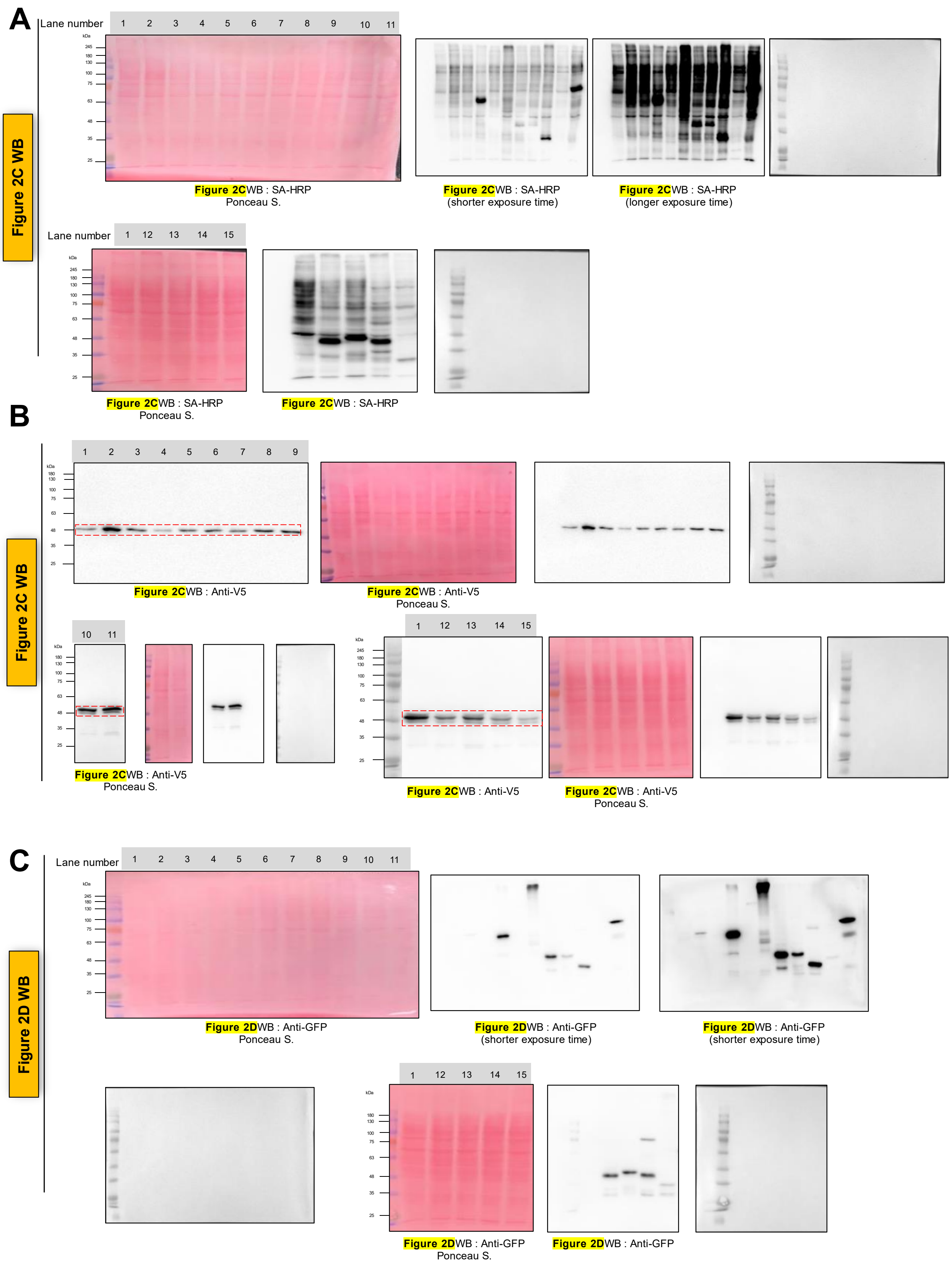


**Supplementary Figure 3. Raw images of western blotting results from Figure 2. (A)** Raw images of western blotting and Ponceau membrane stain results with size ladder from **Figure 2c** (WB: SA-HRP). **(B)** Raw images of western blotting and Ponceau membrane staining results with size ladder from **Figure 2c** (WB: anti-V5). Selected areas for **Figure 2c** are marked with red box. **(C)** Raw images of western blotting and Ponceau membrane staining results with size ladder from **Figure 2d** (WB: anti-GFP).

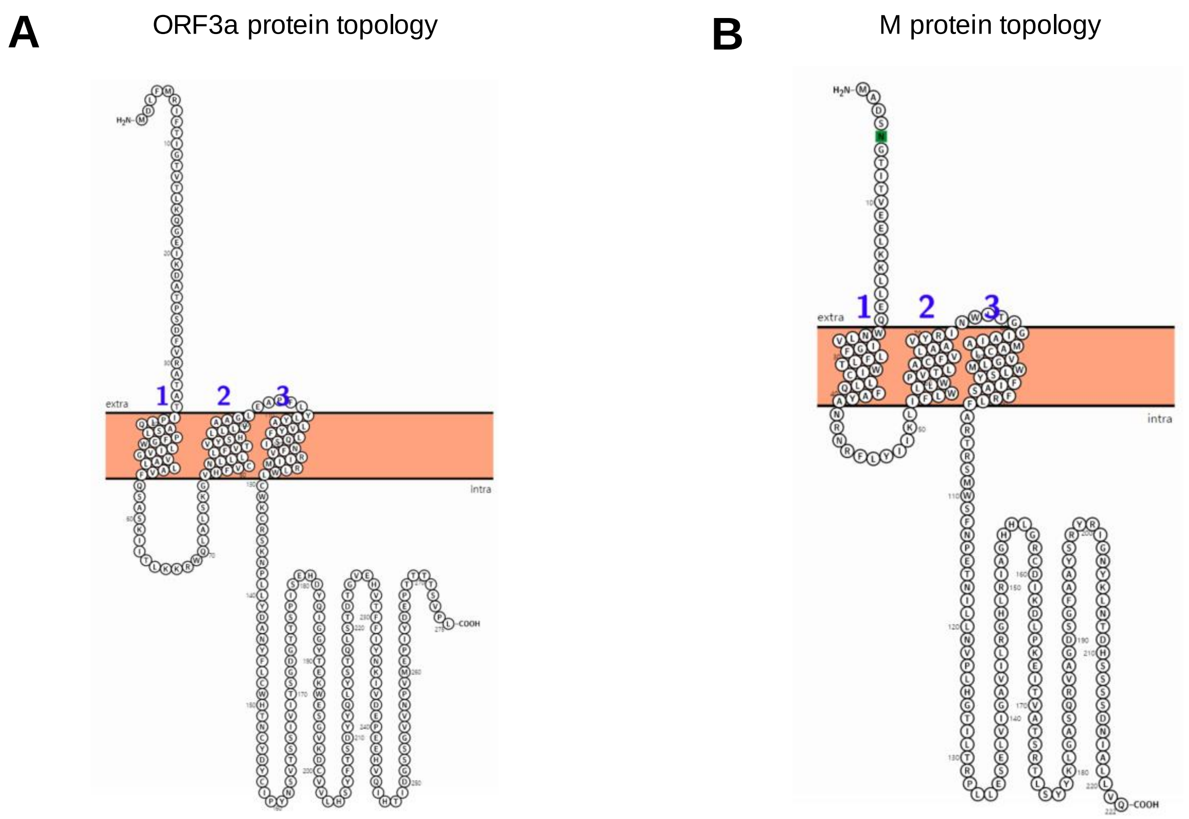


*Protter - visualize proteoforms* *Omasits et al., Bioinformatics. 2013 Nov 21.*

**Supplementary Figure 4. Predicted transmembrane domain regions of ORF3a and M by Protter. (A)** Membrane topology of ORF3a. **(B)** Membrane topology of M These data was obtained by using Protter (https://wlab.ethz.ch/protter/start/). N-glyco motif was shown with green box.

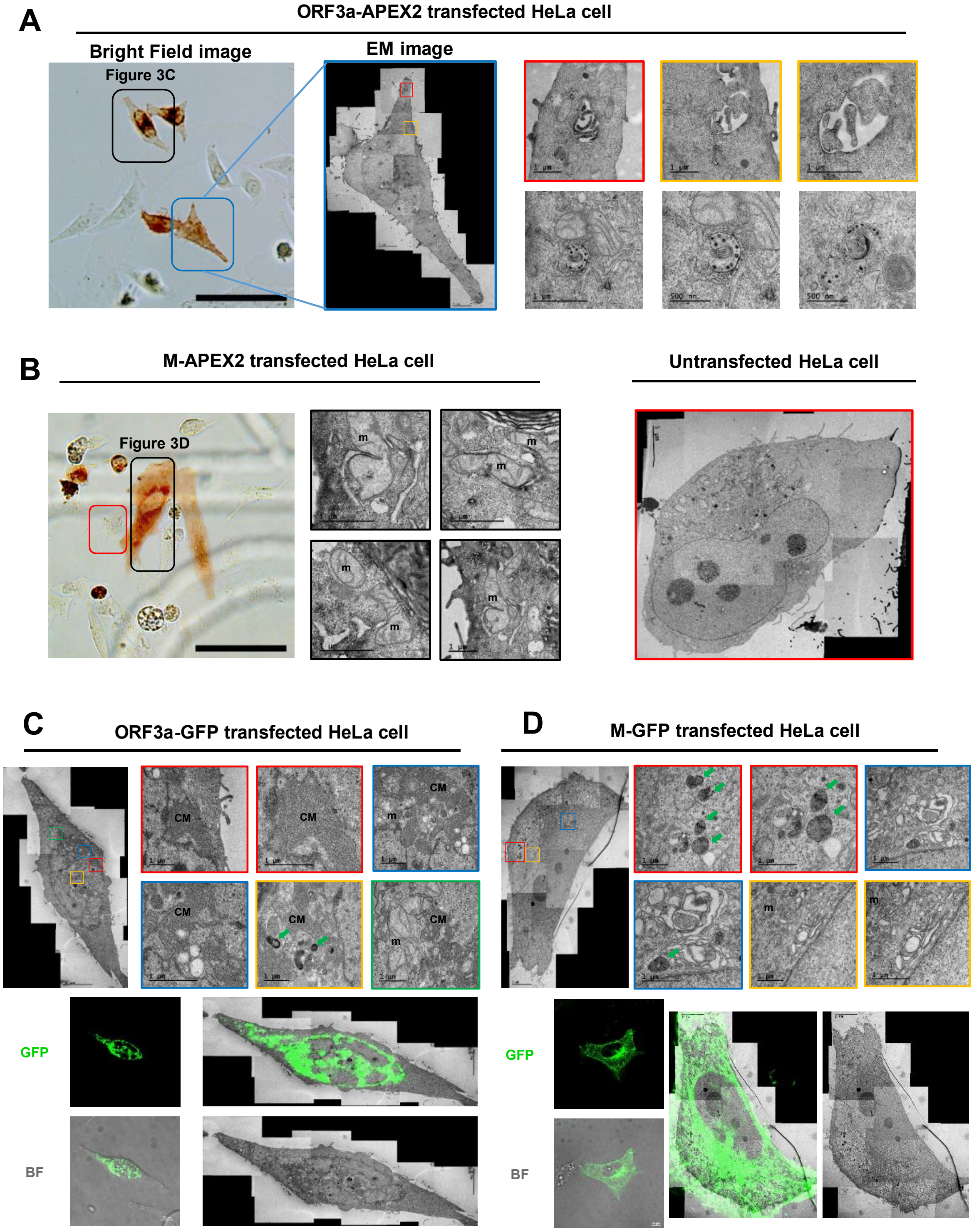


**Supplementary Figure 5. Additional EM images of ORF3a or M expressed HeLa cells. (A)** Additional EM images of ORF3a-linker-V5-APEX2 transfected HeLa cells. Bright field image of DAB-stained ORF3a-APEX2 transfected HeLa cells were shown in left. An EM image of the cell shown in **Figure 3c** is marked with black box in a bright field image. Scale bars : 500 m - 1 μm. **(B)** Additional EM images of M-linker-V5-APEX2 transfected HeLa cells. An EM image of the cell shown in **Figure 3d** is marked with black box in the bright field image. Untransfected cell was also shown with red box. Mitochondria is indicated as "m", respectively (b–d). Scale bars: 1 μm. **(C)** CLEM images of ORF3a-GFP transfected HeLa cells. Scale bars: 1 μm. Distinct membrane structures were observed with higher magnification. Magnified area are marked with various color boxes.) Cubic membranes are indicated as "CM". **(D)** CLEM images of M-GFP transfected HeLa cells. Electron-dense autophagic vesicles are marked with green arrows. Scale bars: 1 μm.

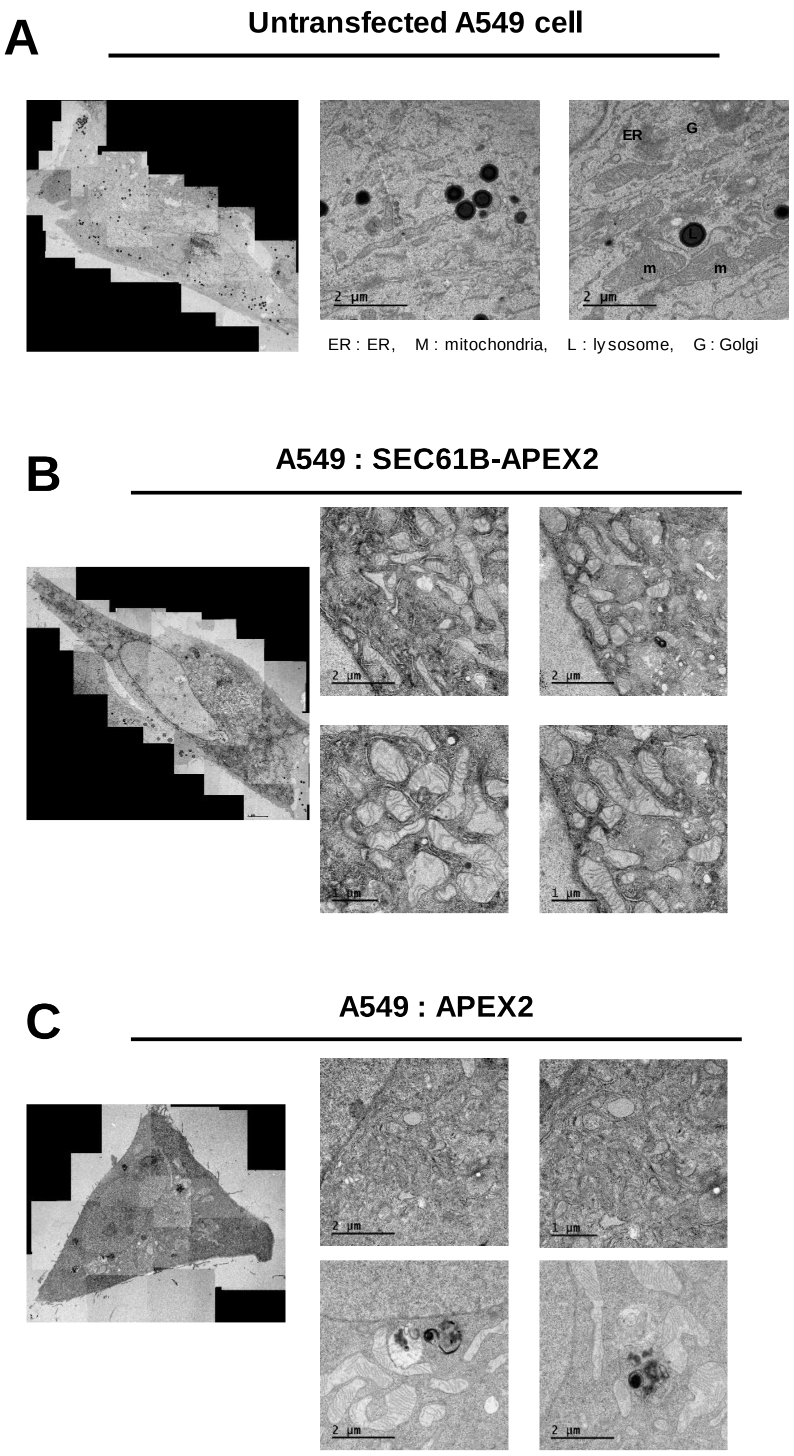


**Supplementary Figure 6. EM images of control samples of A549 cells. (A)** EM images of non-transfected A549 cells. Scale bars: 2 μm. **(B)** EM images of SEC61B-APEX2 transfected A549 cells. Scale bars: 1 - 2 μm. **(C)** EM images of APEX2 transfected cells. Scale bars: 1 - 2 μm.

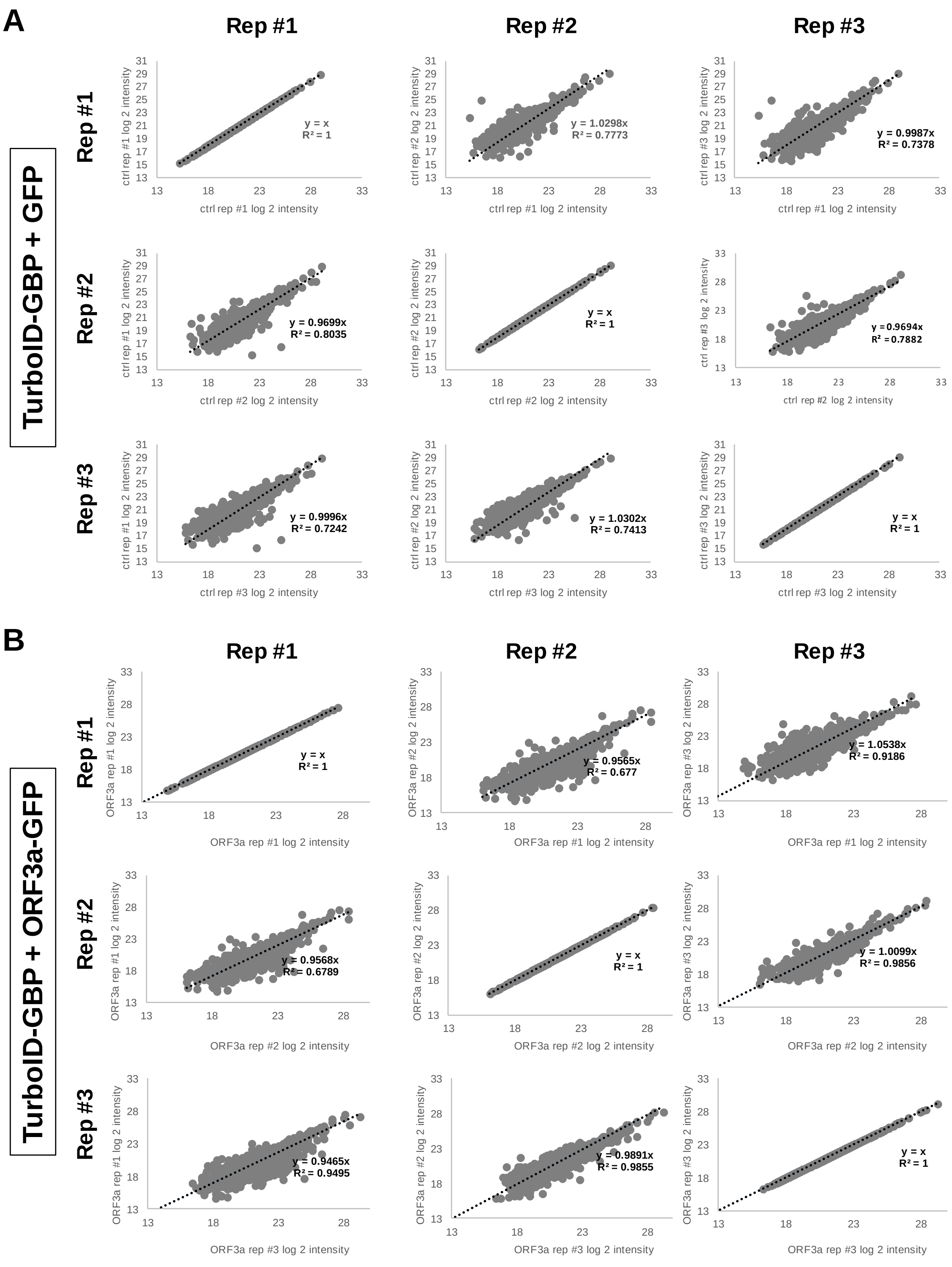
A
Rep #1
Rep #2
Rep #3
TurboID-GBP + GFP
ctrl rep #1 log 2 intensity
ctrl rep #2 log 2 intensity
ctrl rep #3 log 2 intensity
y = x
R² = 1
y = 1.0298x
R² = 0.7773
y = 0.9987x
R² = 0.7378
y = 0.9699x
R² = 0.8035
y = x
R² = 1
y = 0.9694x
R² = 0.7882
y = 0.9996x
R² = 0.7242
y = 1.0302x
R² = 0.7413
y = x
R² = 1
B
Rep #1
Rep #2
Rep #3
TurboID-GBP + ORF3a-GFP
ORF3a rep #1 log 2 intensity
ORF3a rep #2 log 2 intensity
ORF3a rep #3 log 2 intensity
y = x
R² = 1
y = 0.9565x
R² = 0.677
y = 1.0538x
R² = 0.9186
y = 0.9568x
R² = 0.6789
y = x
R² = 1
y = 1.0099x
R² = 0.9856
y = 0.9465x
R² = 0.9495
y = 0.9891x
R² = 0.9855
y = x
R² = 1

**C**

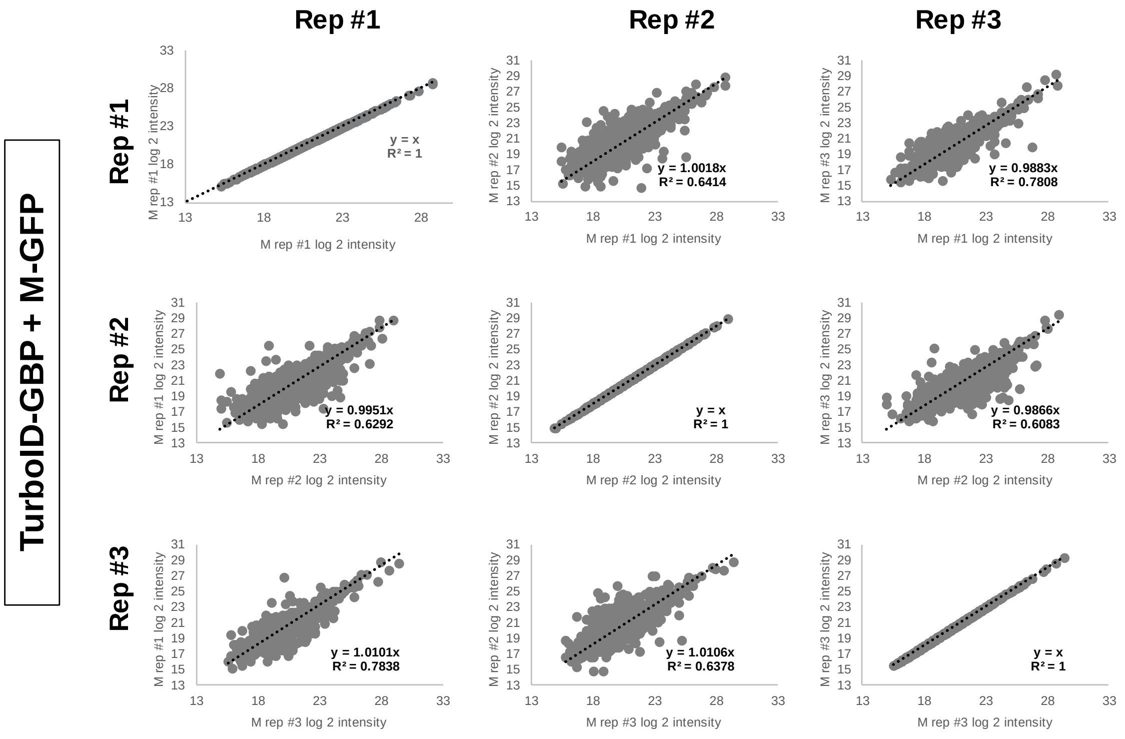


**D**

| | | ctrl : GFP + TurboID-GBP | | | ORF3a-GFP + TurboID-GBP | | | M-GFP + TurboID-GBP | | |
|---|---|---|---|---|---|---|---|---|---|---|
| | | rep #1 | rep #2 | rep #3 | rep #1 | rep #2 | rep #3 | rep #1 | rep #2 | rep #3 |
| ctrl : GFP + TurboID-GBP | rep #1 | 1 | 0.924343 | 0.9441 | 0.484594 | 0.428742 | 0.291283 | 0.266151 | 0.210742 | 0.19369 |
| | rep #2 | 0.924343 | 1 | 0.976369 | 0.581784 | 0.489374 | 0.396407 | 0.325799 | 0.262611 | 0.226385 |
| | rep #3 | 0.9441 | 0.976369 | 1 | 0.559304 | 0.454481 | 0.333395 | 0.294569 | 0.216243 | 0.180331 |
| ORF3a-GFP + TurboID-GBP | rep #1 | 0.484594 | 0.581784 | 0.559304 | 1 | 0.845807 | 0.845415 | 0.869613 | 0.768219 | 0.751899 |
| | rep #2 | 0.428742 | 0.489374 | 0.454481 | 0.845807 | 1 | 0.93272 | 0.780329 | 0.833266 | 0.765891 |
| | rep #3 | 0.291283 | 0.396407 | 0.333395 | 0.845415 | 0.93272 | 1 | 0.853849 | 0.887966 | 0.872773 |
| M-GFP + TurboID-GBP | rep #1 | 0.266151 | 0.325799 | 0.294569 | 0.869613 | 0.780329 | 0.853849 | 1 | 0.881657 | 0.878903 |
| | rep #2 | 0.210742 | 0.262611 | 0.216243 | 0.768219 | 0.833266 | 0.887966 | 0.881657 | 1 | 0.907887 |
| | rep #3 | 0.19369 | 0.226385 | 0.180331 | 0.751899 | 0.765891 | 0.872773 | 0.878903 | 0.907887 | 1 |

Low [color scale] High

**E**

### ORF3a interactome : 117 proteins

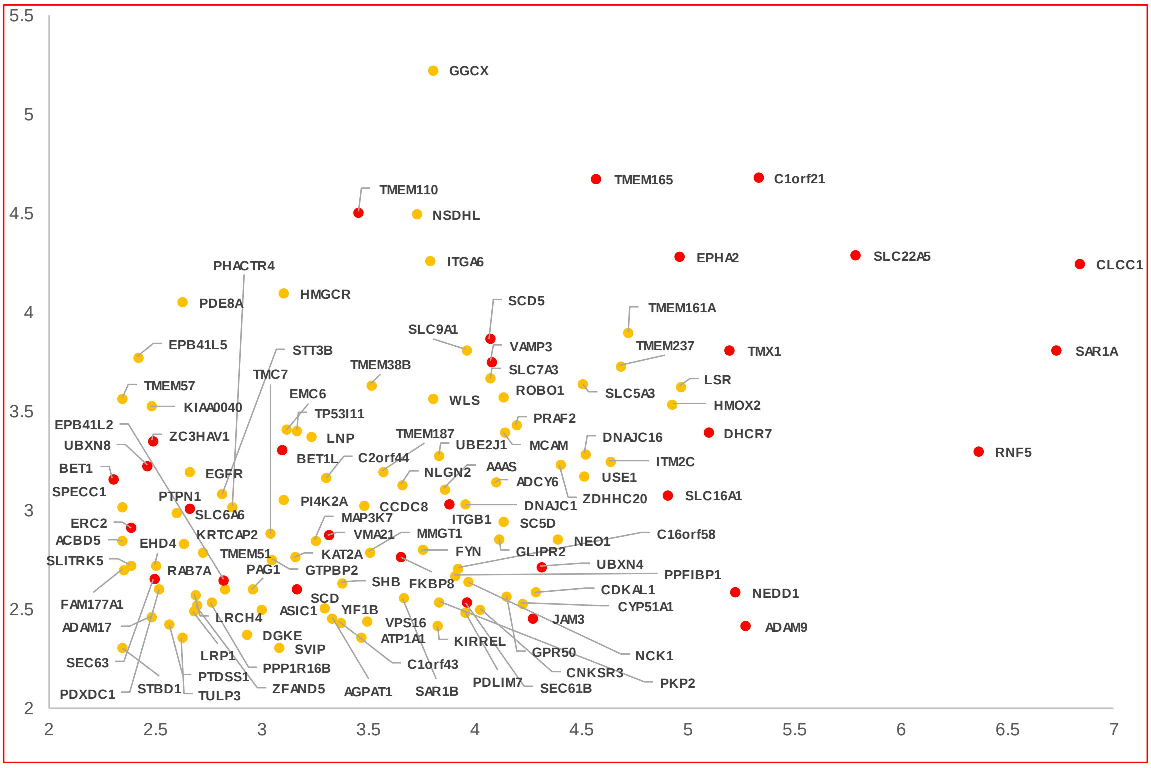


### M interactome : 191 proteins

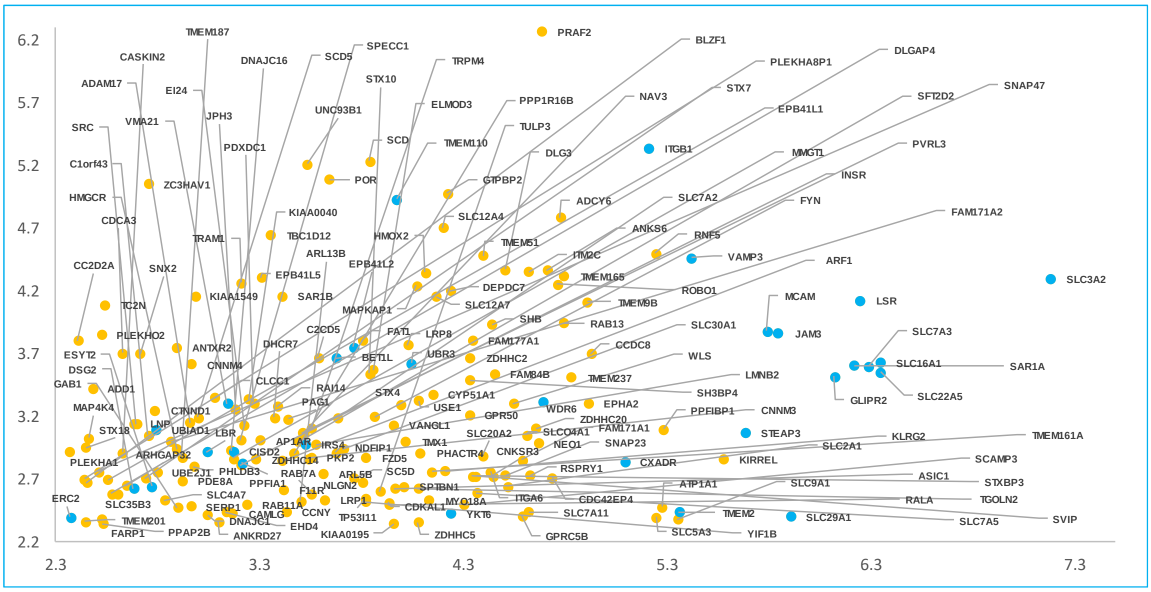

**F**

### ORF3a interactome : 117 proteins

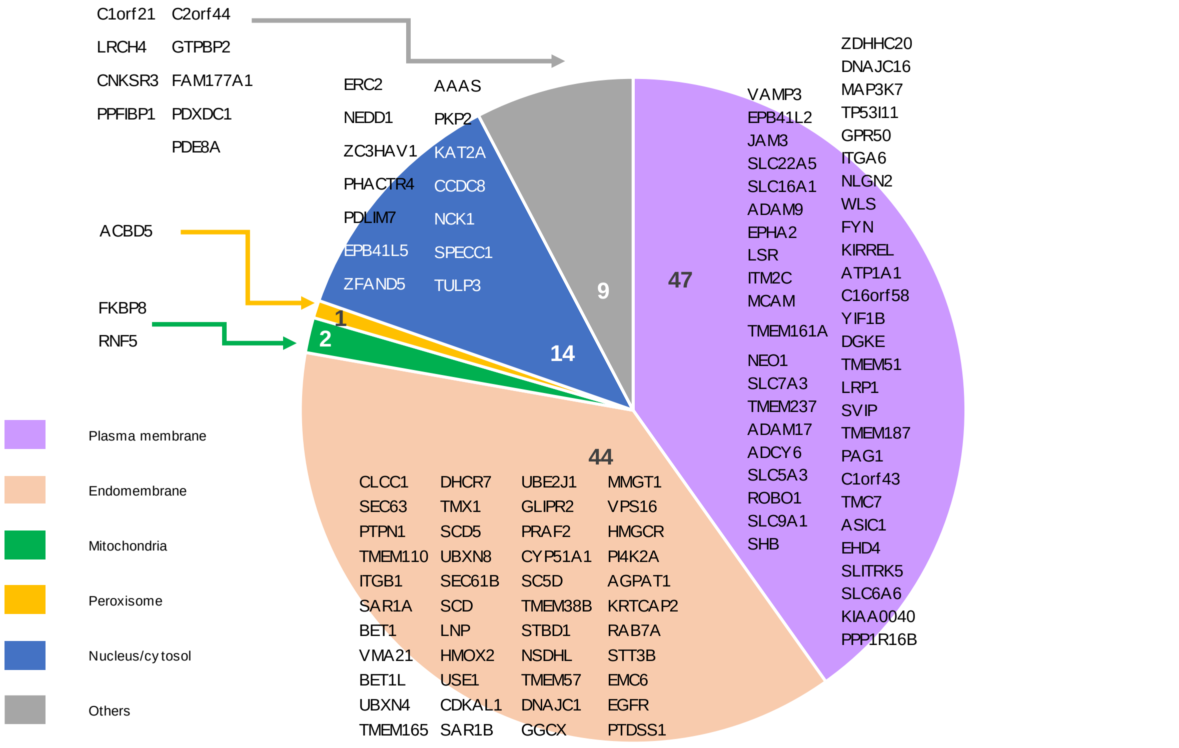


### M interactome : 191 proteins

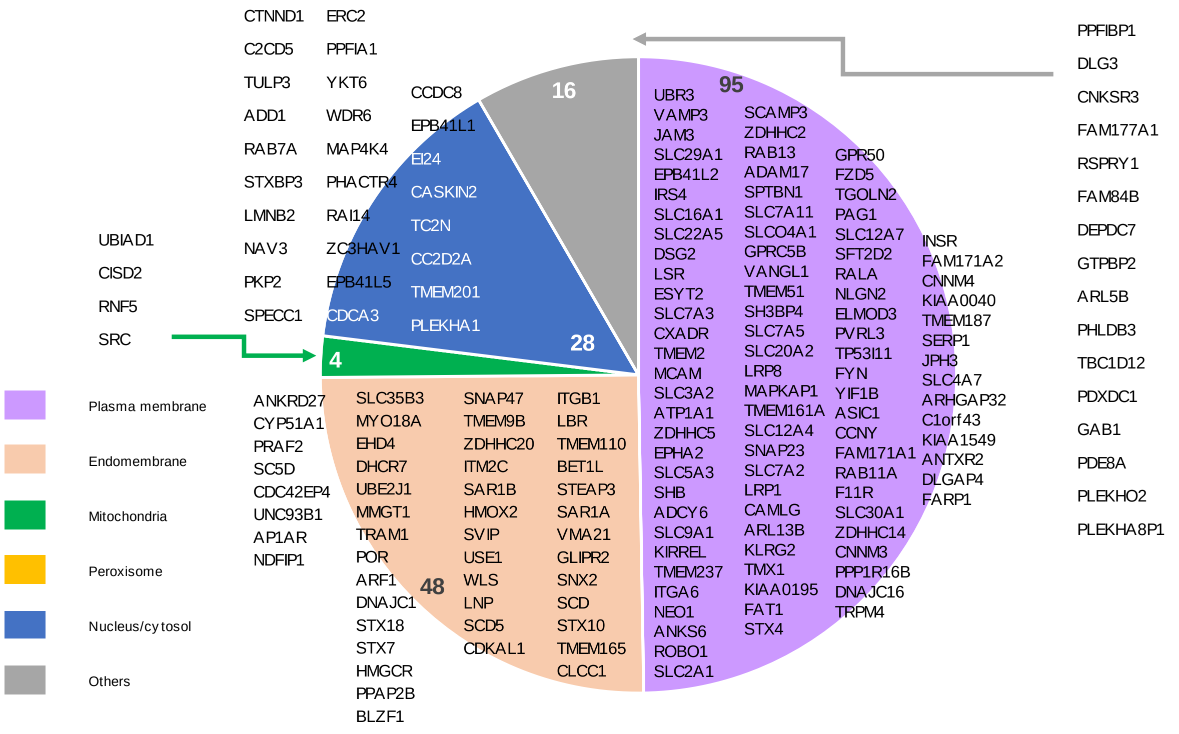

G

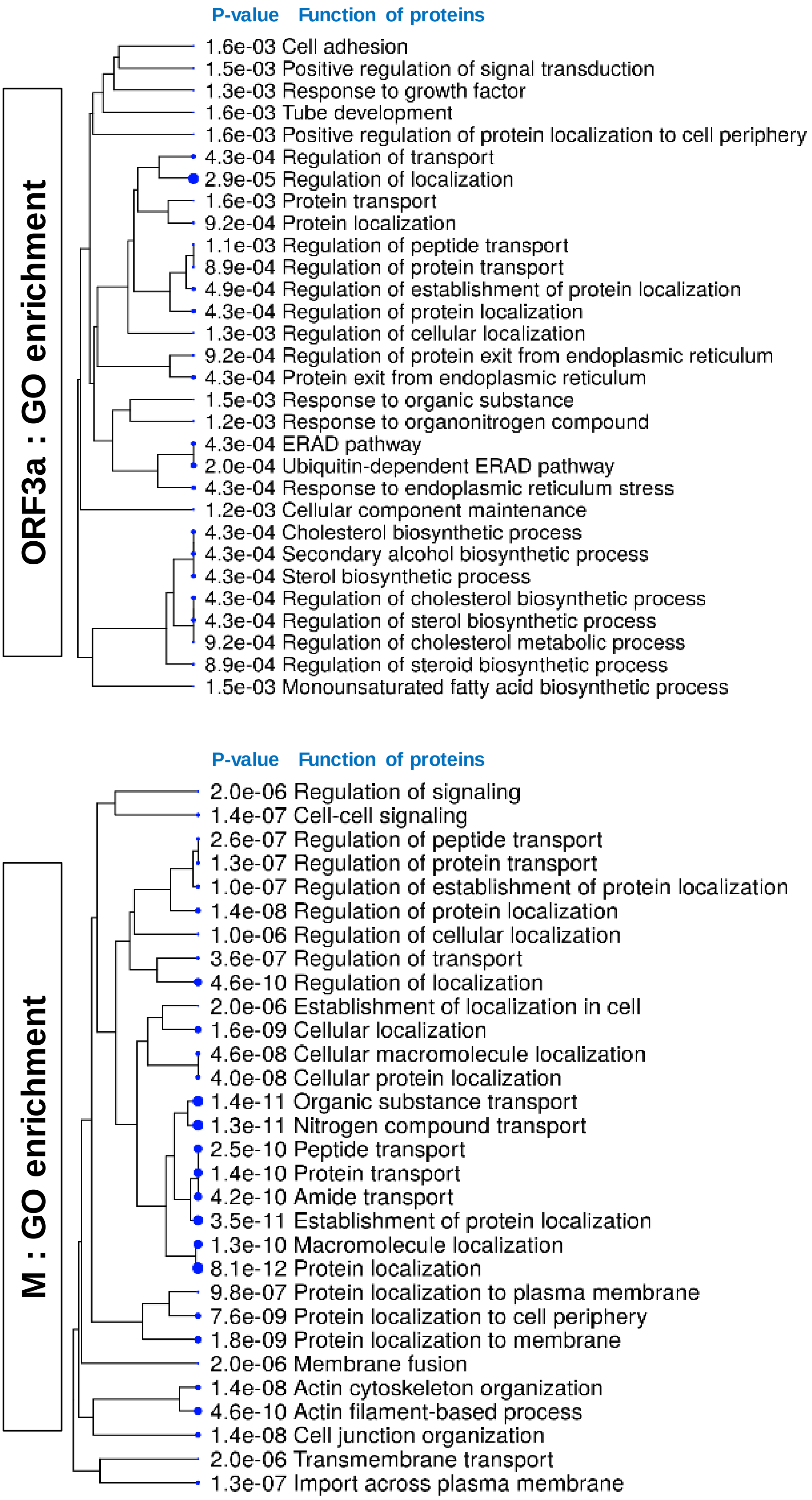
ORF3a : GO enrichment
P-value Function of proteins
1.6e-03 Cell adhesion
1.5e-03 Positive regulation of signal transduction
1.3e-03 Response to growth factor
1.6e-03 Tube development
1.6e-03 Positive regulation of protein localization to cell periphery
4.3e-04 Regulation of transport
2.9e-05 Regulation of localization
1.6e-03 Protein transport
9.2e-04 Protein localization
1.1e-03 Regulation of peptide transport
8.9e-04 Regulation of protein transport
4.9e-04 Regulation of establishment of protein localization
4.3e-04 Regulation of protein localization
1.3e-03 Regulation of cellular localization
9.2e-04 Regulation of protein exit from endoplasmic reticulum
4.3e-04 Protein exit from endoplasmic reticulum
1.5e-03 Response to organic substance
1.2e-03 Response to organonitrogen compound
4.3e-04 ERAD pathway
2.0e-04 Ubiquitin-dependent ERAD pathway
4.3e-04 Response to endoplasmic reticulum stress
1.2e-03 Cellular component maintenance
4.3e-04 Cholesterol biosynthetic process
4.3e-04 Secondary alcohol biosynthetic process
4.3e-04 Sterol biosynthetic process
4.3e-04 Regulation of cholesterol biosynthetic process
4.3e-04 Regulation of sterol biosynthetic process
9.2e-04 Regulation of cholesterol metabolic process
8.9e-04 Regulation of steroid biosynthetic process
1.5e-03 Monounsaturated fatty acid biosynthetic process
M : GO enrichment
P-value Function of proteins
2.0e-06 Regulation of signaling
1.4e-07 Cell-cell signaling
2.6e-07 Regulation of peptide transport
1.3e-07 Regulation of protein transport
1.0e-07 Regulation of establishment of protein localization
1.4e-08 Regulation of protein localization
1.0e-06 Regulation of cellular localization
3.6e-07 Regulation of transport
4.6e-10 Regulation of localization
2.0e-06 Establishment of localization in cell
1.6e-09 Cellular localization
4.6e-08 Cellular macromolecule localization
4.0e-08 Cellular protein localization
1.4e-11 Organic substance transport
1.3e-11 Nitrogen compound transport
2.5e-10 Peptide transport
1.4e-10 Protein transport
4.2e-10 Amide transport
3.5e-11 Establishment of protein localization
1.3e-10 Macromolecule localization
8.1e-12 Protein localization
9.8e-07 Protein localization to plasma membrane
7.6e-09 Protein localization to cell periphery
1.8e-09 Protein localization to membrane
2.0e-06 Membrane fusion
1.4e-08 Actin cytoskeleton organization
4.6e-10 Actin filament-based process
1.4e-08 Cell junction organization
2.0e-06 Transmembrane transport
1.3e-07 Import across plasma membrane

**Supplementary Figure 7. Additional information of mass analysis results (A)** Scatter plot of MS1 intensity of biotinylated peptides from biological triplicate samples ofGFP (control, i.e. TurboID-GBP and GFP). **(B)** Scatter plot of MS1 intensity of biotinylated peptides from biological triplicate samples of ORF3a-GFP (i.e. TurboID-GBP and ORF3a-GFP). **(C)** Scatter plot of MS1 intensity of biotinylated peptides from biological triplicate samples of M-GFP (i.e. TurboID-GBP and M-GFP). **(D)** Correlation value table of MS1 intensities of biotinylated peptides in biological tripicates samples of GFP, ORF3a-GFP and M-GFP **(E)** Expended view of ORF3a and M interactomes with gene names in the volcano plot of **Figure 4b**. 117 proteins at ORF3a interactome and 191 proteins at M interactome were all shown. Detail information is shown in Supplementary Dataset 1. **(F)** Subcellular distribution of ORF3a and M interactome shown in **Figure 4c**. **(G)** Functional enrichment of ORF3a interactome (117 proteins) and M interactome (191 proteins). Gene ontology analysis was performed using ShinyGO v0.61: Gene Ontology Enrichment Analysis (http://bioinformatics.sdstate.edu/go/).

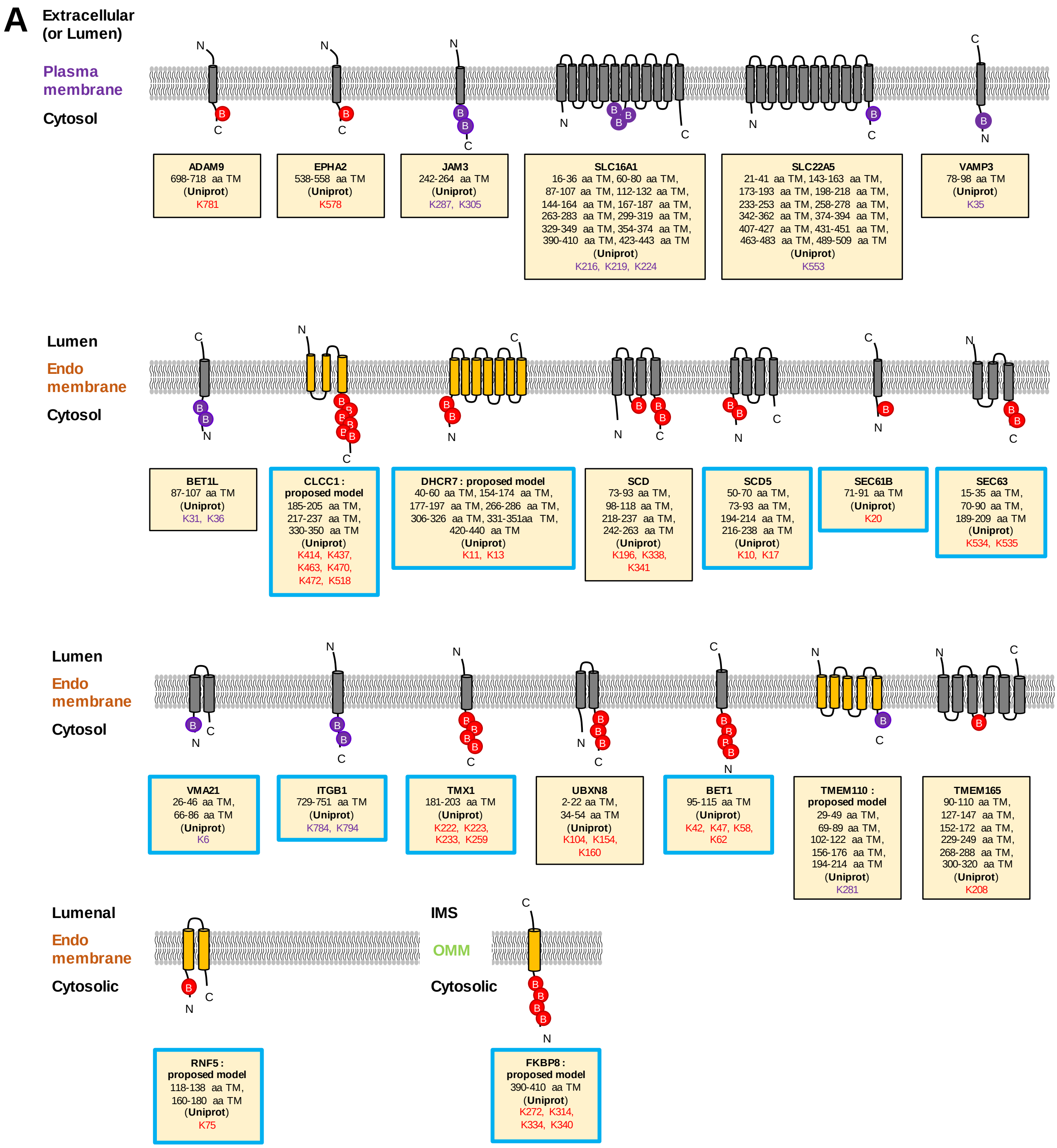
A
Extracellular (or Lumen)
Plasma membrane
Cytosol
ADAM9 698-718 aa TM (Uniprot) K781
EPHA2 538-558 aa TM (Uniprot) K578
JAM3 242-264 aa TM (Uniprot) K287, K305
SLC16A1 16-36 aa TM, 60-80 aa TM, 87-107 aa TM, 112-132 aa TM, 144-164 aa TM, 167-187 aa TM, 263-283 aa TM, 299-319 aa TM, 329-349 aa TM, 354-374 aa TM, 390-410 aa TM, 423-443 aa TM (Uniprot) K216, K219, K224
SLC22A5 21-41 aa TM, 143-163 aa TM, 173-193 aa TM, 198-218 aa TM, 233-253 aa TM, 258-278 aa TM, 342-362 aa TM, 374-394 aa TM, 407-427 aa TM, 431-451 aa TM, 463-483 aa TM, 489-509 aa TM (Uniprot) K553
VAMP3 78-98 aa TM (Uniprot) K35
Lumen
Endo membrane
Cytosol
BET1L 87-107 aa TM (Uniprot) K31, K36
CLCC1 : proposed model 185-205 aa TM, 217-237 aa TM, 330-350 aa TM (Uniprot) K414, K437, K463, K470, K472, K518
DHCR7 : proposed model 40-60 aa TM, 154-174 aa TM, 177-197 aa TM, 266-286 aa TM, 306-326 aa TM, 331-351aa TM, 420-440 aa TM (Uniprot) K11, K13
SCD 73-93 aa TM, 98-118 aa TM, 218-237 aa TM, 242-263 aa TM (Uniprot) K196, K338, K341
SCD5 50-70 aa TM, 73-93 aa TM, 194-214 aa TM, 216-238 aa TM (Uniprot) K10, K17
SEC61B 71-91 aa TM (Uniprot) K20
SEC63 15-35 aa TM, 70-90 aa TM, 189-209 aa TM (Uniprot) K534, K535
Lumen
Endo membrane
Cytosol
VMA21 26-46 aa TM, 66-86 aa TM (Uniprot) K6
ITGB1 729-751 aa TM (Uniprot) K784, K794
TMX1 181-203 aa TM (Uniprot) K222, K223, K233, K259
UBXN8 2-22 aa TM, 34-54 aa TM (Uniprot) K104, K154, K160
BET1 95-115 aa TM (Uniprot) K42, K47, K58, K62
TMEM110 : proposed model 29-49 aa TM, 69-89 aa TM, 102-122 aa TM, 156-176 aa TM, 194-214 aa TM (Uniprot) K281
TMEM165 90-110 aa TM, 127-147 aa TM, 152-172 aa TM, 229-249 aa TM, 268-288 aa TM, 300-320 aa TM (Uniprot) K208
Lumenal
Endo membrane
Cytosolic
RNF5 : proposed model 118-138 aa TM, 160-180 aa TM (Uniprot) K75
IMS
OMM
Cytosolic
FKBP8 : proposed model 390-410 aa TM (Uniprot) K272, K314, K334, K340

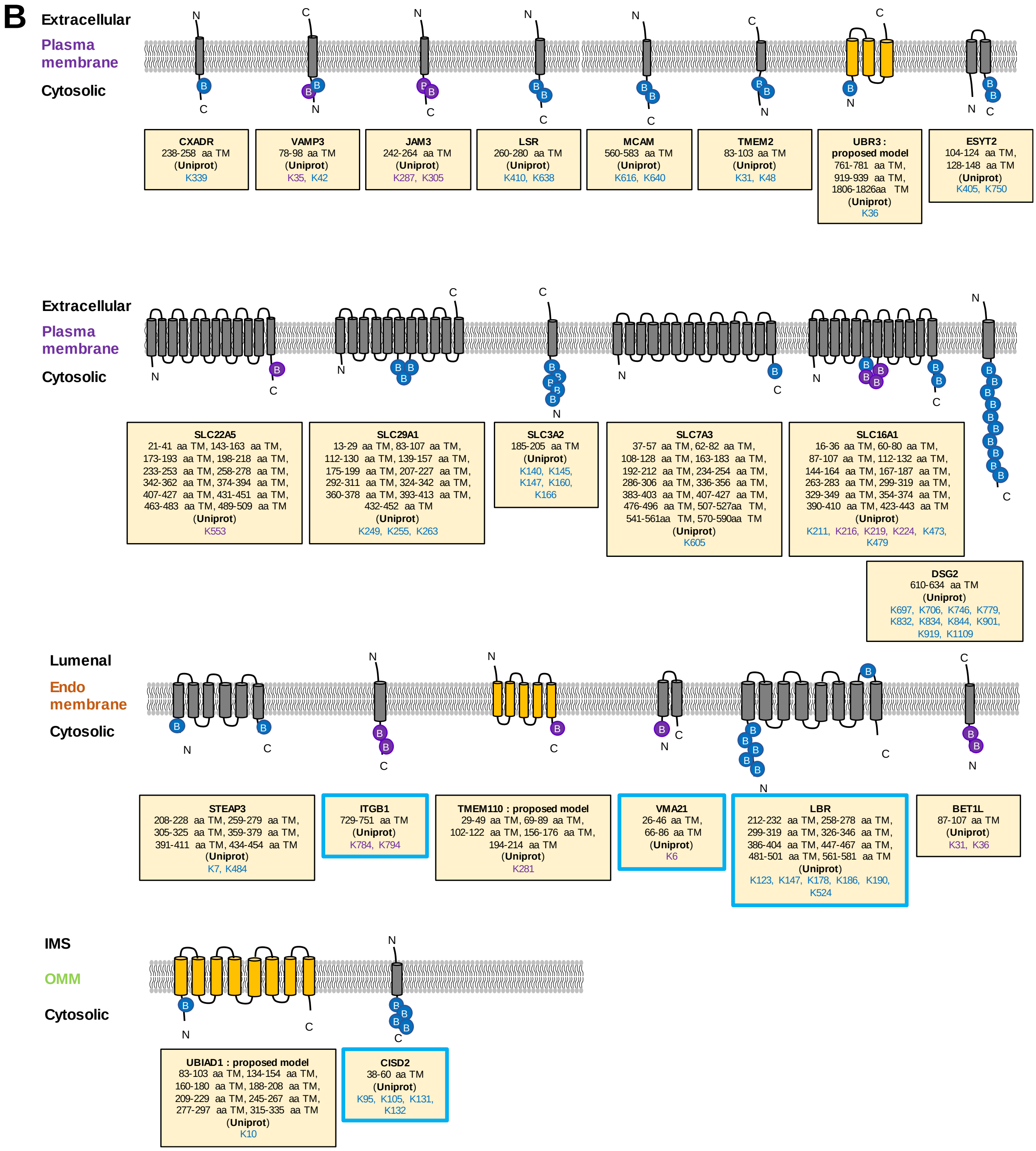


**Supplementary Figure 8. Proposed membrane topology of host membrane proteins of ORF3a and M of SARS-CoV-2 from Figure 5. (A)** Results of biotin-labeled sites on membrane protein by ORF3a-GFP and TurboID-GBP. Among 30 proteins at ORF3a interactome of **Figure 4e**, 22 proteins are integral membrane proteins. Biotin-labeled sites in ORF3a-GFP sample, not in M-GFP sample are shown as red-colored biotin ("B"). Biotin-labeled sites in both ORF3a- and M-GFP samples are shown as purple-colored biotin. Proposed models of topology (e.g. CLCC1, DHCR7, TMEM110, FKBP8, RNF5) based on our mass data were colored in yellow. Among depicted 22 proteins, proteins overlapped with MAM proteome were shown with light blue colored boxes. IMS meant mitochondrial intermembrane space. OMM meant outer mitochondrial membrane. **(B)** Results of biotin-labeled sites on membrane protein by M-GFP and TurboID-GBP. Among 30 proteins at M interactome of **Figure 4f**, 22 proteins are integral membrane proteins. Biotin-labeled sites by M not ORF3a are shown as blue-colored biotin. Biotin-labeled sites by both ORF3a and M are shown as purple-colored biotin. Proposed models of topology (e.g. UBR3, TMEM110, UBIAD1) based on our mass data were colored in yellow. Among depicted 22 proteins, proteins overlapped with MAM proteome were shown with light blue colored boxes. IMS meant mitochondrial intermembrane space. OMM meant outer mitochondrial membrane.

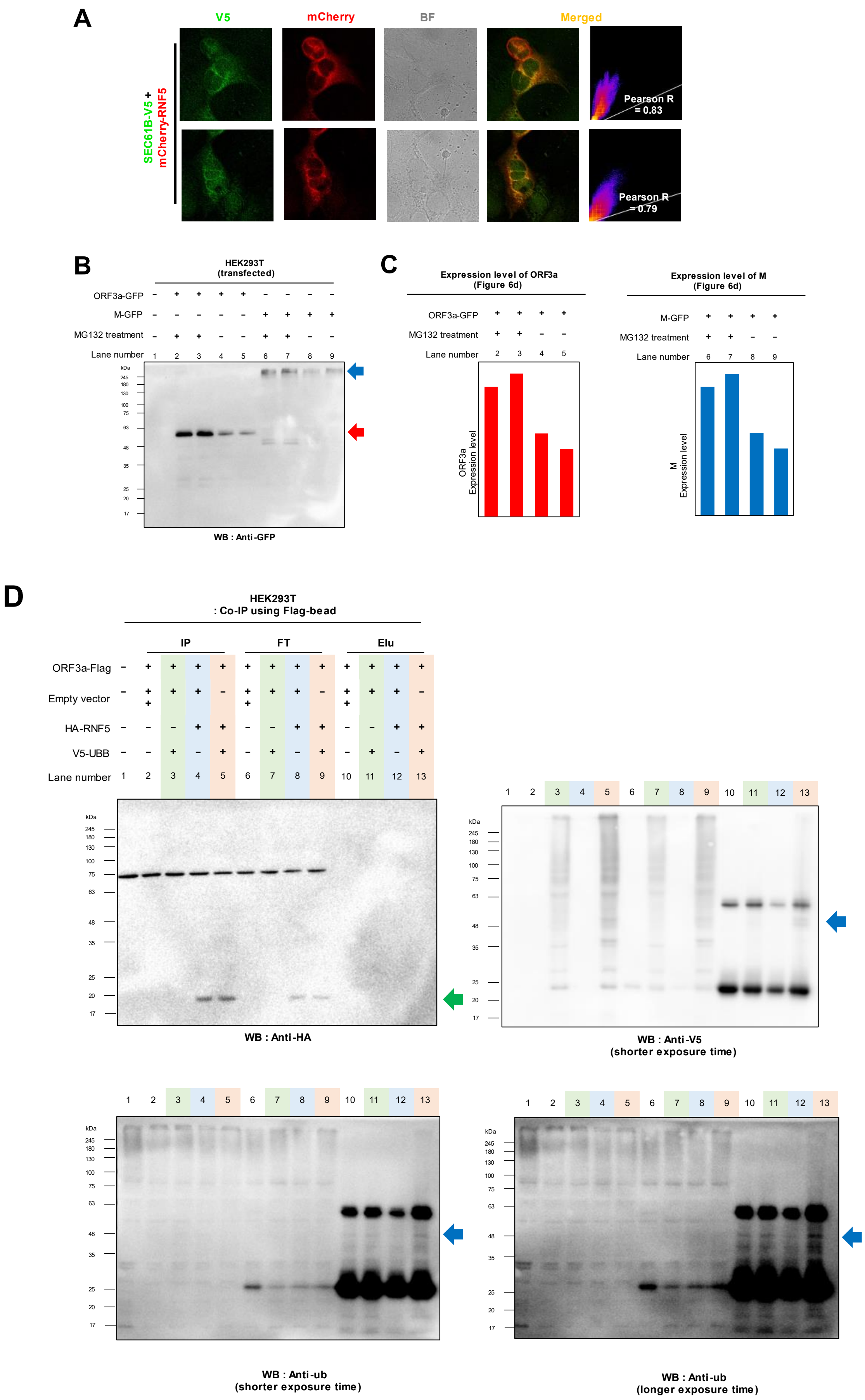
A
V5
mCherry
BF
Merged
SEC61B-V5 +
mCherry-RNF5
Pearson R = 0.83
Pearson R = 0.79
B
HEK293T
(transfected)
ORF3a-GFP
M-GFP
MG132 treatment
Lane number
kDa
WB : Anti-GFP
C
Expression level of ORF3a
(Figure 6d)
ORF3a
Expression level
Expression level of M
(Figure 6d)
M
Expression level
D
HEK293T
: Co-IP using Flag-bead
IP
FT
Elu
ORF3a-Flag
Empty vector
HA-RNF5
V5-UBB
Lane number
WB : Anti-HA
WB : Anti-V5
(shorter exposure time)
WB : Anti-ub
(shorter exposure time)
WB : Anti-ub
(longer exposure time)

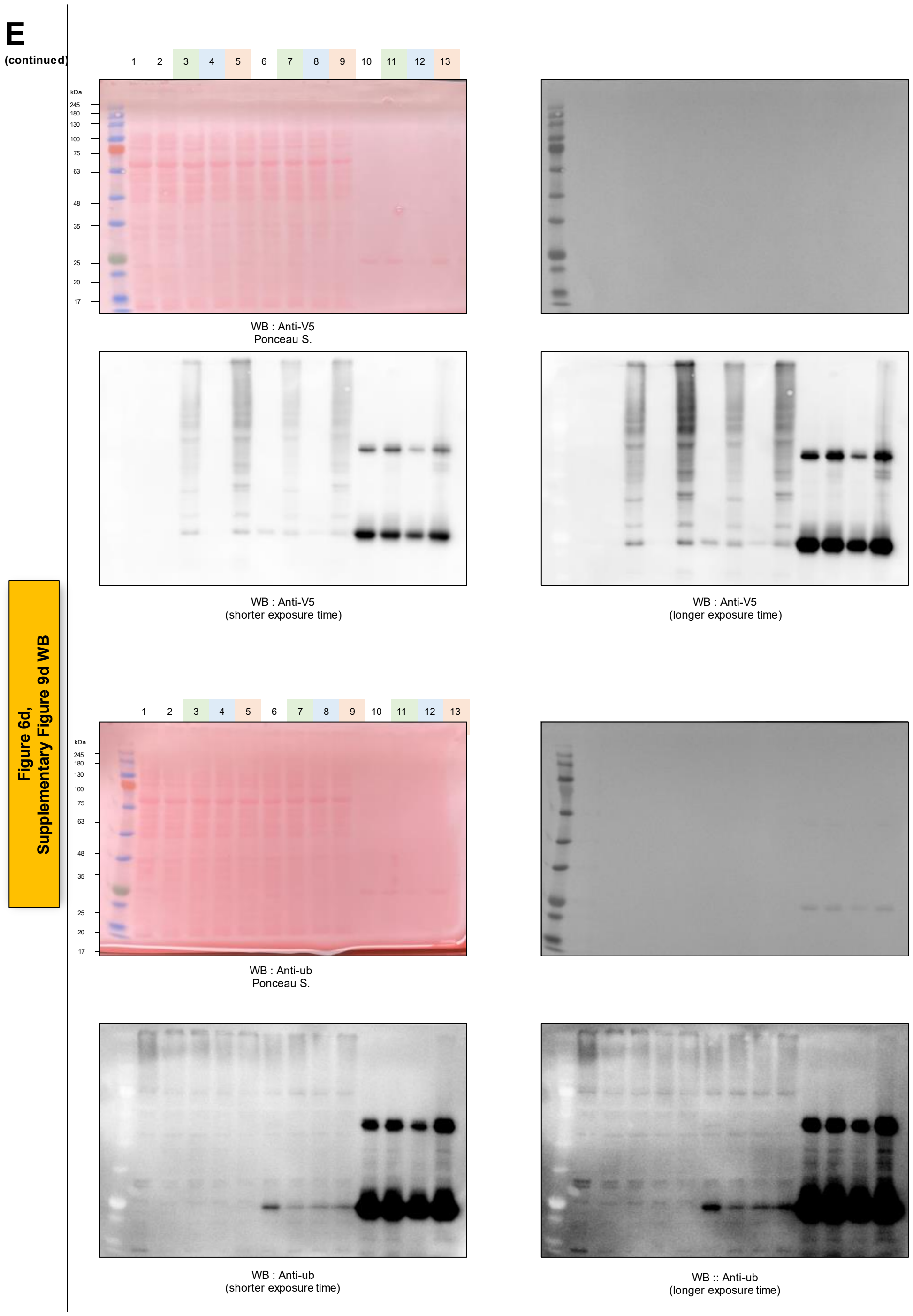


**Supplementary Figure 9. RNF5 ubiquitinylates ORF3a of SARS-CoV-2. (a)** Confocal microscopy images of Flag-RNF5 (left) and SEC61-V5-TurboID with mCherry-RNF5. Flag-RNF5 was visualized by anti-Flag antibody (AF488-conjugated, GFP fluorescence channel). SEC61B-V5-TurboID was visualized by anti-V5 antibody (AF488-conjugated, GFP fluorescence channel). mCherry-RNF5 was visualized by RFP fluorescence channel. BF: bright field, Scale bars: 10 µm. **(b)** Anti-GFP Western blot results of ORF3a-GFP or M-GFP expressed HEK293T cells with or without MG132 treatment (2 µM, 16 h). **(c)** Expression of ORF3a and M shown in (b). After MG132 treatment, expression of ORF3a and M increased. **(d)** Additional western blotting results are shown in **Figure 6d**. Anti-HA Anti-V5, and Anti-ub western blotting results of ORF3a-Flag, HA-RNF5 and V5-UBB co-expressed HEK293T cells. Band with red arrow is the expected size of ORF3a-Flag. Band with blue arrows is the ubiquitinylated ORF3a-Flag. Band with green arrow is HA-RNF5. **(e)** Raw images of western blotting and Ponceau membrane staining with size ladder from (d) (WB: Anti-Flag, Anti-HA, Anti-V5 and Anti-ub).

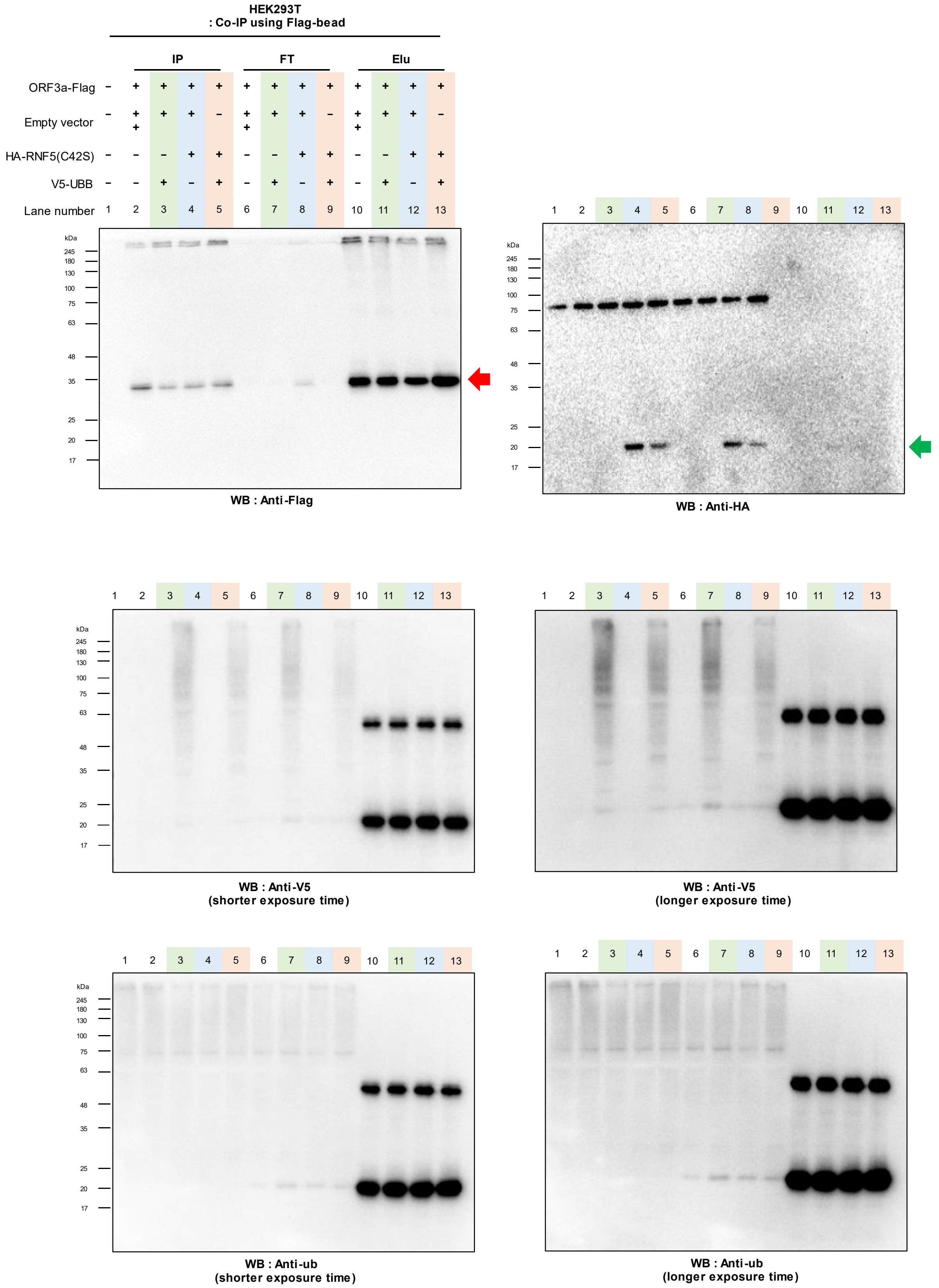


**Supplementary Figure 10. RNF5$^{C42S}$ mutant do not induce ubiquitination of SARS-CoV-2 ORF3a** Anti-Flag, Anti-HA, Anti-V5 and Anti-ubiquitin (ub) western blot results of ORF3a-Flag, HA-RNF5(C42S) and V5-UBB co-expressed HEK293T cells. Red arrow marked unmodified molecular weight of ORF3a-Flag. Band with green arrow is HA-RNF5.

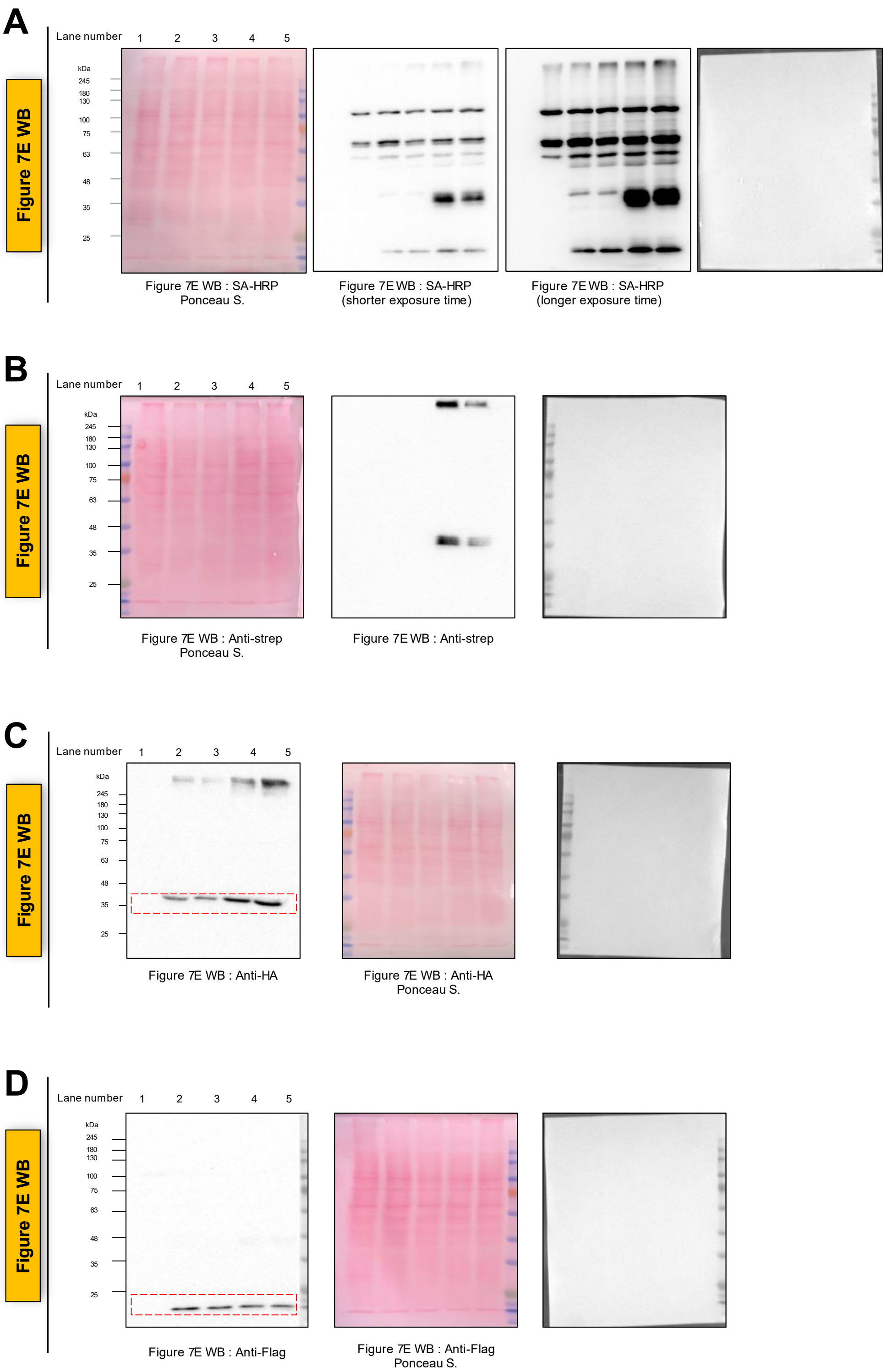


**Supplementary Figure 11. Raw images of western blot results from Figure 7e. (A)** Raw images of western blot and ponceau membrane staining results with size ladder of **Figure 7e** (WB : SA-HRP). **(B)** Raw images of western blot and ponceau membrane staining results with size ladder of **Figure 7e** (WB : anti-strep). **(C)** Raw images of western blot and ponceau membrane staining results with size ladder of **Figure 7e** (WB : anti-HA). Parts surrounded with red dotted line were shown at **Figure 7e**. **(D)** Raw images of western blot and ponceau membrane staining results with size ladder of **Figure 7e** (WB : anti-FLAG). Parts surrounded with red dotted line were shown at **Figure 7e**.

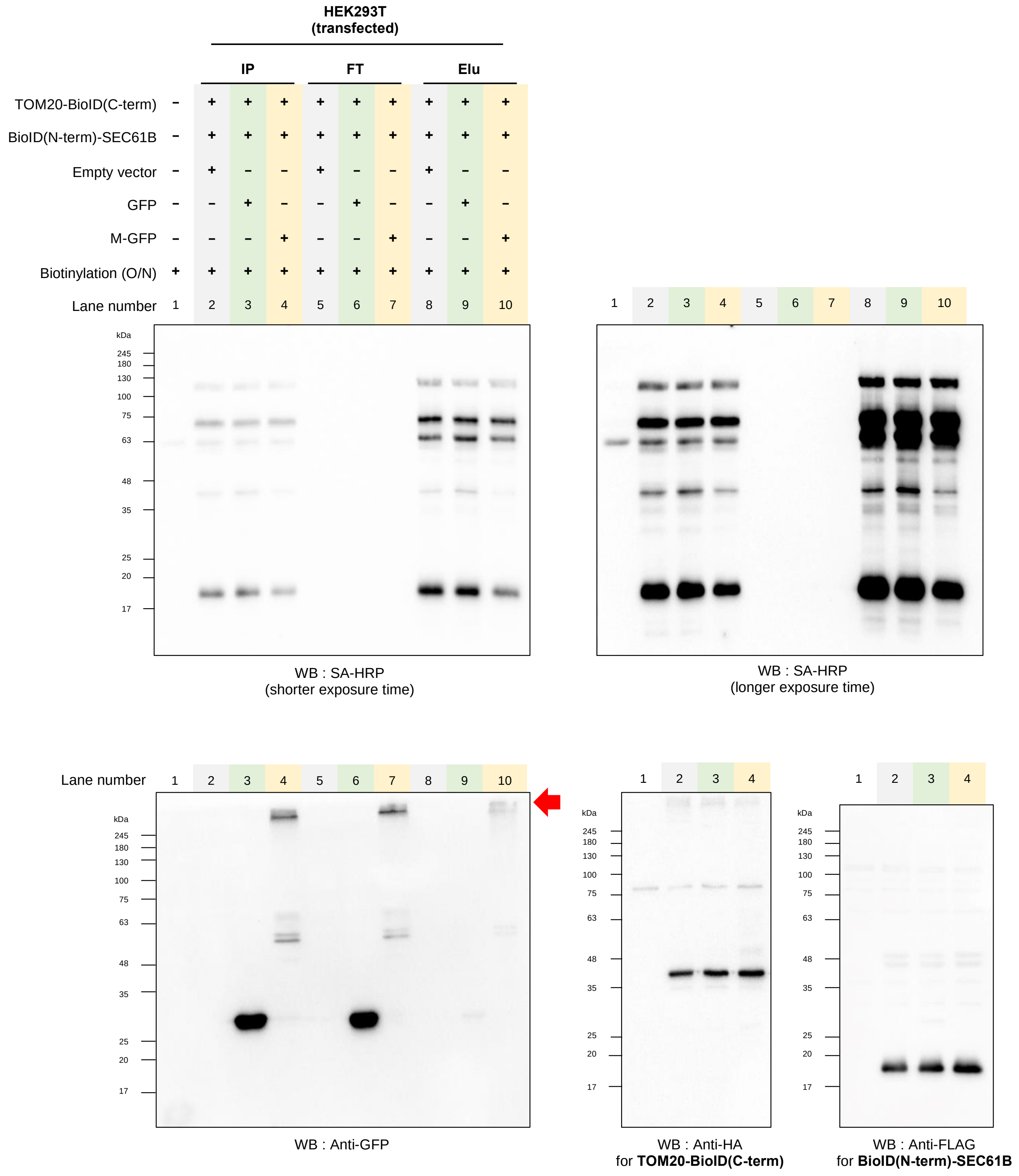


**Supplementary Figure 12. Western blot results of M expressed cells using Contact-ID.** SA-HRP, Anti-GFP, Anti-HA and Anti-Flag western blot results of whole cell lysate of GFP and M-GFP expressed HEK293T cells with co-expression of Contact-ID constructs. These samples were enriched by SA-bead. M was biotinylated by Contact-ID constructs, indicating M localized to MAM. IP, FT and Elu meant input, flow through and elution of samples. Bands with red arrow showed M protein biotinylated by Contact-ID.

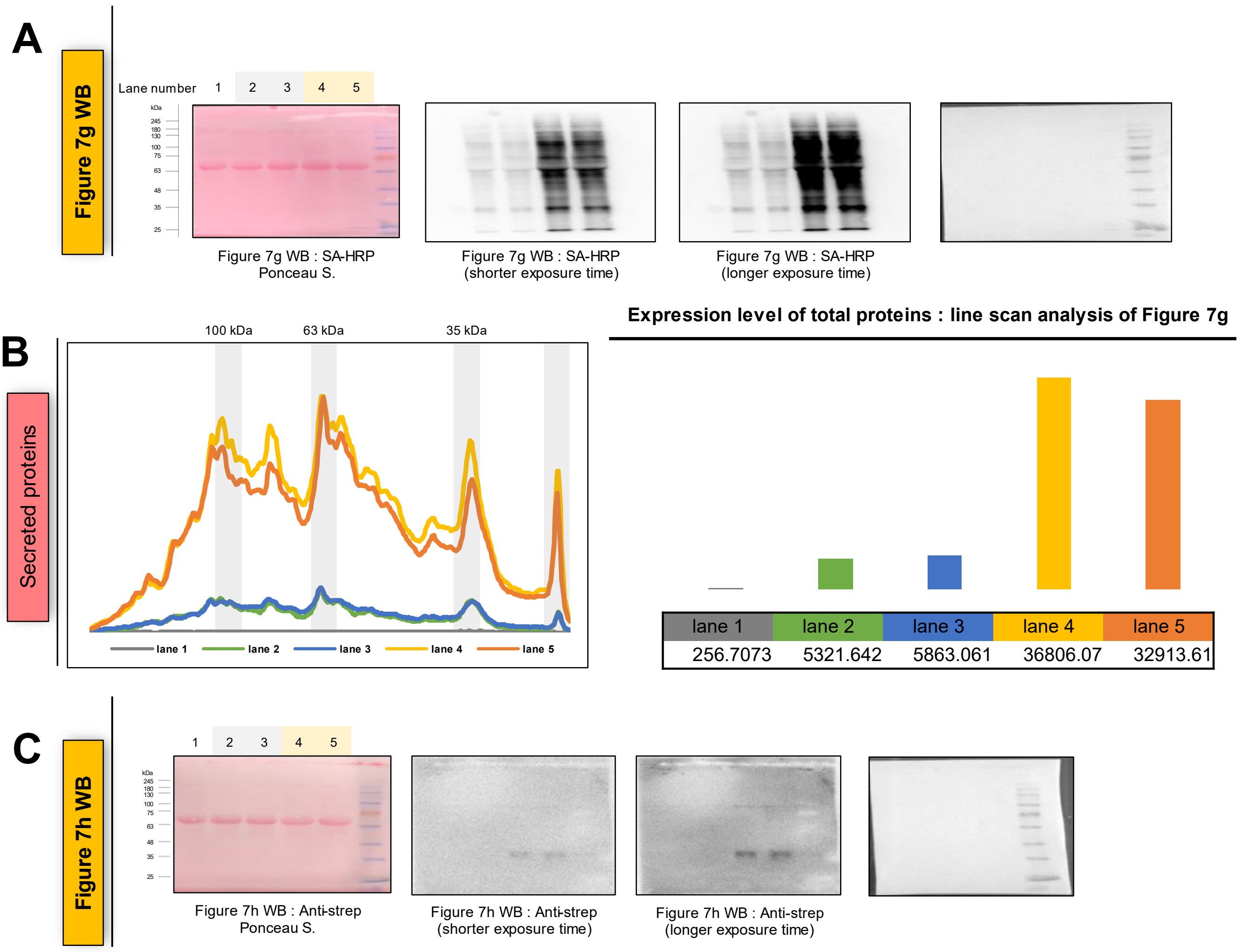


**Supplementary Figure 13. ORF3a affects secretome profiles of host cells. (A)** Raw images of western blotting and Ponceau membrane staining with size ladder from **Figure 7g** (WB: SA-HRP). **(B)** Line scan analysis results of **Figure 7g**. Expression of proteins from **Figure 7g** is shown in detail. Total expression of secreted proteins from **Figure 7g**. From densitometric analysis of the western blotting lanes, we observed that an increase in biotinylated proteins of over 6.23 folds was detected in ORF3a-expressing cells. **(C)** Raw images of western blotting and Ponceau membrane staining with size ladder from **Figure 7h** (WB: anti-strep).

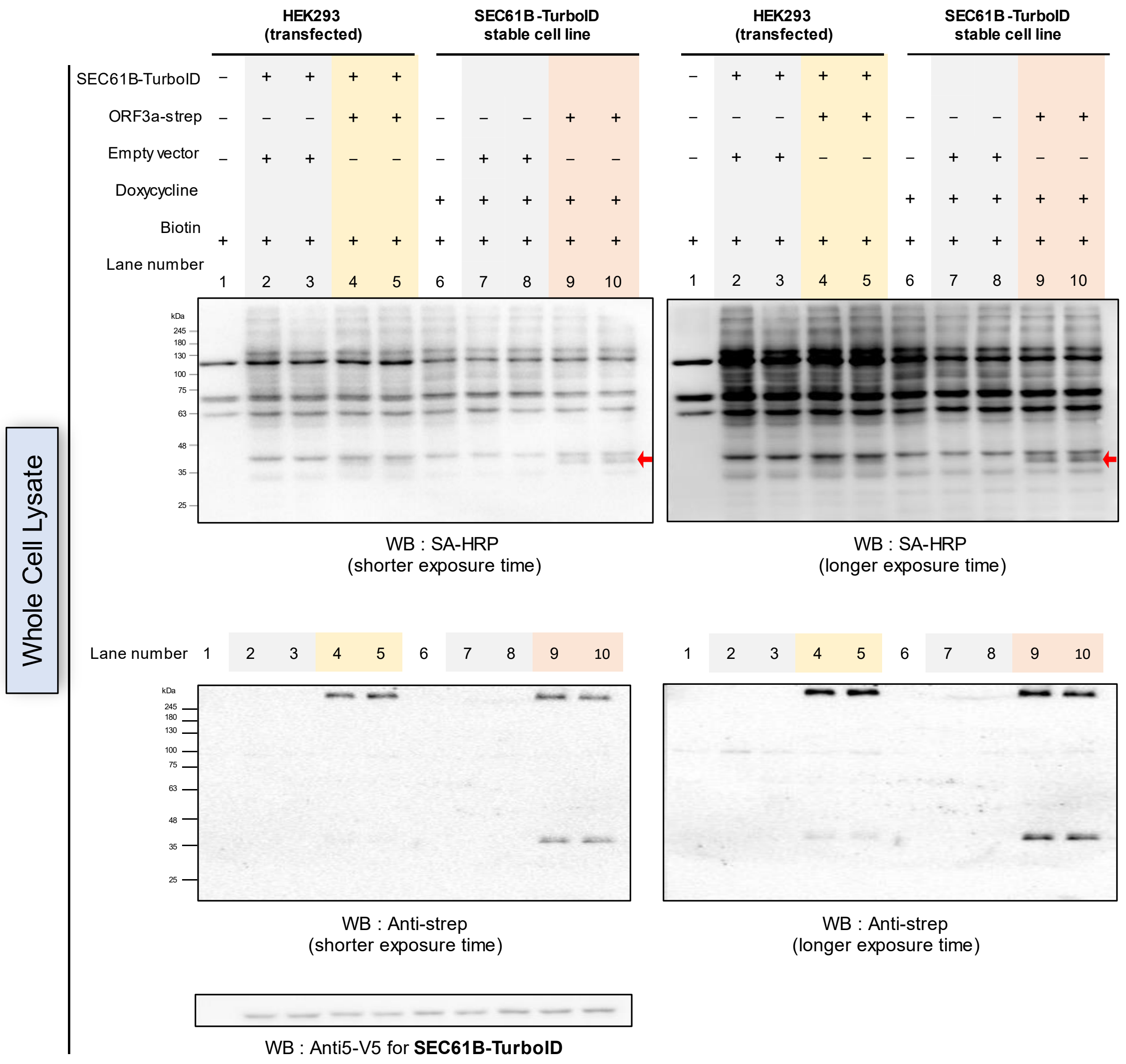


**Supplementary Figure 14. Western blot results of whole cell lysates under ORF3a expression using iSLET method.** SA-HRP, Anti-strep and Anti-V5 western blot results of whole cell lysate of SEC61B-TurboID expressed HEK293 cells with or without co-expression of ORF3a-strep. Bands of biotinylated ORF3a-strep (in at lane 4,5 and 9,10) were marked with red arrows. The biotinylating activity of SEC61B-TurboID was not affected by the addition of ORF3a expression since the total amount of biotinylated proteins in whole cell lysates was rarely affected by ORF3a expression, and this was evident in cells with transient and stable expression of SEC61B-TurboID.

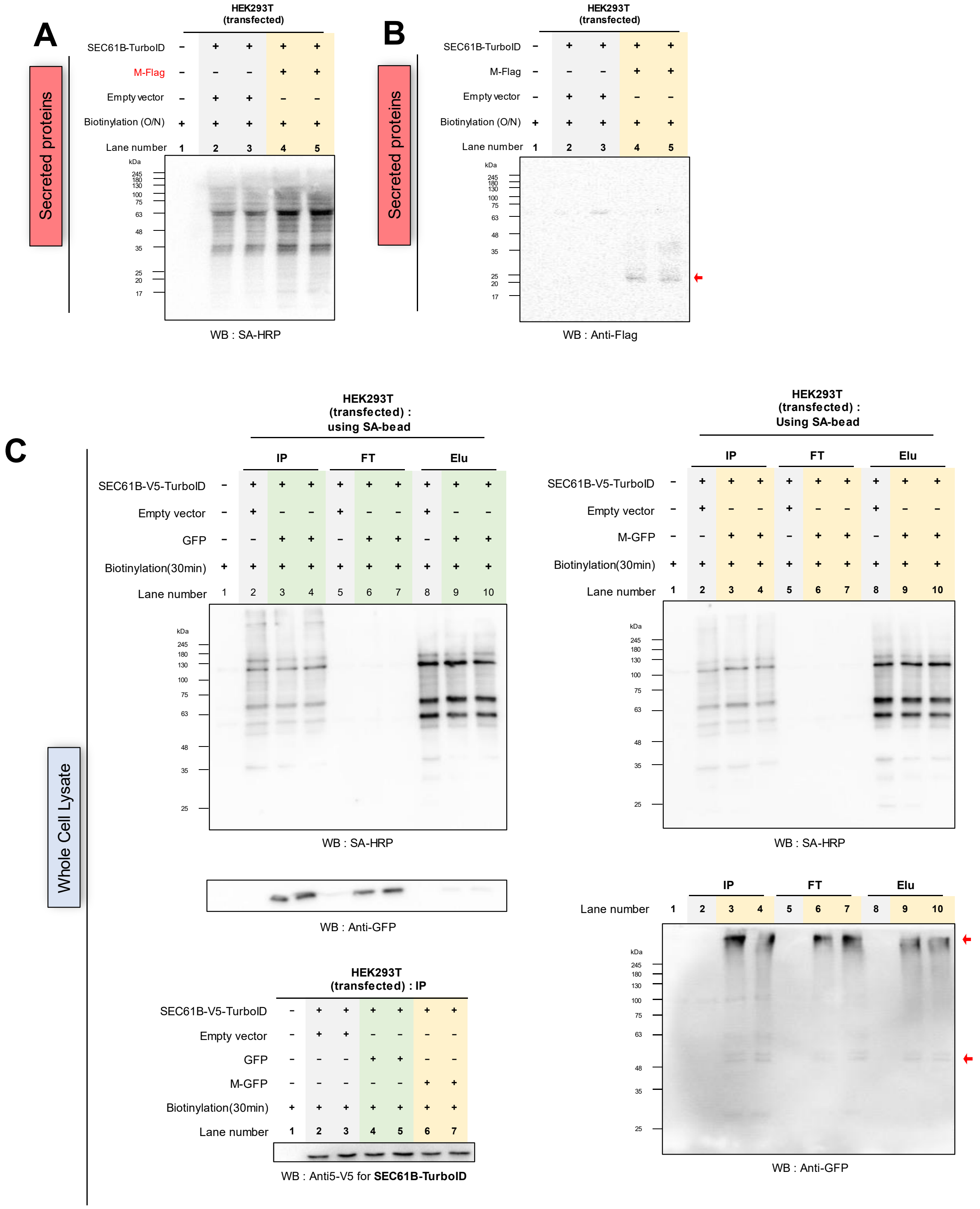


**Supplementary Figure 15. M affects secretome profiles of host cells. (a)** SA-HRP western blotting result of secreted proteins from SEC61B-TurboID expressed HEK293T cells with or without co-expression of M. **(b)** Anti-Flag western blotting results of the same samples as **(a)**. **(c)** Whole cell lysate sample from **(a)**.

**Supplementary Table 1. Construct Information**

Templates of nsp7, nsp8, nsp9, nsp16, S, E, M, N, ORF3a, ORF6, ORF7a, ORF7b, and ORF8 were obtained from professor Kim, Ho Min (KAIST). Templates of ORF3b, ORF9b, ORF9c, and ORF10 were obtained from professor Kim, V. Narry (SNU).

| Name (expected size) | Features | Promotor/Vector | Details |
|---|---|---|---|
| nsp7-linker-eGFP (expected MW : 37.7 kDa) | *HindIII*-*KpnI*-nsp7-*BsiWI*-linker-*NheI*-eGFP-Stop-*NotI*-*XhoI* | CMV/pCDNA5 | linker : GAPGSAGSAAGSG |
| nsp8-linker-eGFP (expected MW : 50.3 kDa) | *HindIII*-*KpnI*-nsp8-*BsiWI*-linker-*NheI*-eGFP-Stop-*NotI*-*XhoI* | CMV/pCDNA5 | linker : GAPGSAGSAAGSG |
| nsp9-linker-eGFP (expected MW : 40.7 kDa) | *AflII*-nsp9-*HindIII*-linker-*NotI*-eGFP-Stop-*AsCI*-*XhoI* | CMV/pCDNA5 | linker : GAPGSAGSAAGSG |
| nsp16-linker-eGFP (expected MW : 61.6 kDa) | *AflII*-nsp16-*HindIII*-linker-*NotI*-eGFP-Stop-*AsCI*-*XhoI* | CMV/pCDNA5 | linker : GAPGSAGSAAGSG |
| S-linker-eGFP (expected MW : 169.5 kDa) | *HindIII*-*KpnI*-Spike-*BsiWI*-linker-*NheI*-eGFP-Stop-*NotI*-*XhoI* | CMV/pCDNA5 | linker : GAPGSAGSAAGSG |
| E-linker-eGFP (expected MW : 36.5 kDa) | *AflII*-Envelope-*HindIII*-linker-*NotI*-eGFP-Stop-*AsCI*-*XhoI* | CMV/pCDNA5 | linker : GAPGSAGSAAGSG |
| M-linker-eGFP (expected MW : 53.3 kDa) | *AflII*-Membrane-*HindIII*-linker-*NotI*-eGFP-Stop-*AsCI*-*XhoI* | CMV/pCDNA5 | linker : GAPGSAGSAAGSG |
| M-Flag (expected MW : 27.6 kDa) | *AflII*-Membrane-*HindIII*-Flag-Stop-*XhoI* | CMV/pCDNA5 | |
| M-linker-mCherry (expected MW : 53.9 kDa) | *AflII*-Membrane-*HindIII*-linker-*NotI*-mCherry-Stop-*AsCI*-*XhoI* | CMV/pCDNA5 | linker : GAPGSAGSAAGSG |
| N-linker-eGFP (expected MW : 73.9 kDa) | *HindIII*-*KpnI*-Nucleocapsid-*BsiWI*-linker-*NheI*-eGFP-Stop-*NotI*-*XhoI* | CMV/pCDNA5 | linker : GAPGSAGSAAGSG |
| ORF3a-linker-eGFP (expected MW : 59.3 kDa) | *AflII*-ORF3a-*HindIII*-linker-*NotI*-eGFP-Stop-*AsCI*-*XhoI* | CMV/pCDNA5 | linker : GAPGSAGSAAGSG |
| ORF3a-Flag (expected MW : 33.6 kDa) | *AflII*-ORF3a-*HindIII*-Flag-Stop-*AsCI*-*XhoI* | CMV/pCDNA5 | |
| ORF3b-linker-eGFP (expected MW : 37.9 kDa) | *AflII*-ORF3b-*BamHI*-*HindIII*-linker-*NotI*-eGFP-Stop-*AsCI*-*XhoI* | CMV/pCDNA5 | linker : GAPGSAGSAAGSG |
| ORF6-linker-eGFP (expected MW : 35.4 kDa) | *AflII*-ORF6-*HindIII*-linker-*NotI*-eGFP-Stop-*AsCI*-*XhoI* | CMV/pCDNA5 | linker : GAPGSAGSAAGSG |
| ORF6-Flag (expected MW : 9.8 kDa) | *AflII*-ORF6-*HindIII*-Flag-Stop-*XhoI* | CMV/pCDNA5 | |
| ORF7a-linker-eGFP (expected MW : 41.9 kDa) | *AflII*-ORF7a-*HindIII*-linker-*NotI*-eGFP-Stop-*AsCI*-*XhoI* | CMV/pCDNA5 | linker : GAPGSAGSAAGSG |
| ORF7b-linker-eGFP (expected MW : 33.5 kDa) | *HindIII*-*KpnI*-ORF7b-*BsiWI*-linker-*NheI*-eGFP-Stop-*NotI*-*XhoI* | CMV/pCDNA5 | linker : GAPGSAGSAAGSG |
| ORF7b-Flag (expected MW : 7.7 kDa) | *HindIII*-*KpnI*-ORF7b-*BsiWI*-Flag-Stop-*XhoI* | CMV/pCNDA5 | |
| ORF8-linker-eGFP | *AflII*-ORF8-*HindIII*- | CMV/pCDNA5 | linker : |

| | | | |
|---|---|---|---|
| (expected MW : 42.0 kDa) | Flag-*NotI*-eGFP-Stop-*AsCI-XhoI* | | GAPGSAGSAAGSG |
| ORF9b-linker-eGFP (expected MW : 39.6 kDa) | *AflII*-ORF9b-*BamHI-HindIII*-linker-*NotI*-eGFP-Stop-*AsCI-XhoI* | CMV/pCDNA5 | linker : GAPGSAGSAAGSG |
| ORF9c-linker-eGFP (expected MW : 39.5 kDa) | *AflII*-ORF9c-*BamHI-HindIII*-linker-*NotI*-eGFP-Stop-*AsCI-XhoI* | CMV/pCDNA5 | linker : GAPGSAGSAAGSG |
| ORF10-linker-eGFP (expected MW : 33.2 kDa) | *AflII*-ORF10-*BamHI-HindIII*-linker-*NotI*-eGFP-Stop-*AsCI-XhoI* | CMV/pCDNA5 | linker : GAPGSAGSAAGSG |
| G3BP1-V5-APEX2-AP (expected MW : 82.6 kDa) | *HindIII-KpnI*-G3BP1-*NheI*-V5-APEX2-AP-Stop-*NotI-XhoI* | CMV/pCDNA5 | for confocal imaging with co-expression of Nucleocapsid |
| eGFP (expected MW : 26.9 kDa) | *HindIII-KpnI*-eGFP-Stop-*NotI-XhoI* | CMV/pCDNA5 | for mass sampling as control |
| TurboID-V5-GBP (expected MW : 49.4 kDa) | *HindIII*-TurboID-*KpnI*-V5-*NotI*-GBP-Stop-*XhoI* | CMV/pCDNA5 | |
| ORF3a-linker-V5-APEX2 (expected MW : 62.6 kDa) | *AflII*-ORF3a-*HindIII*-linker-*NotI*-V5-APEX2-AP-Stop-*XhoI* | CMV/pCDNA5 | linker : GAPGSAGSAAGSG for APEX-EM imaging of ORF3a expression |
| M-linker-V5-APEX2 (expected MW : 56.6 kDa) | *AflII*-Membrane-*HindIII*-linker-*NotI*-V5-APEX2-AP-Stop-*XhoI* | CMV/pCDNA5 | linker : GAPGSAGSAAGSG for APEX-EM imaging of M expression |
| Flag-RNF5 (expected MW : 22.3 kDa) | *AflII*-Flag-RNF5-Stop-*XhoI* | CMV/pCDNA5 | For co-imaging with vPOI |
| mCherry-RNF5 (expected MW : 47.3 kDa) | *AflII*-mCherry-*EcoRI-BamHI*-RNF5-Stop-*XhoI* | CMV/pCNDA5 | For co-imaging with SEC61B |
| HA-RNF5 (expected MW : 21.2 kDa) | *HindIII*-HA-*BamHI*-RNF5-Stop-*XhoI* | CMV/pCNDA5 | For ubiquitination of vPOI |
| HA-RNF5(C42S) (expected MW : 21.2 kDa) | *HindIII*-HA-*BamHI*-RNF5(C42S)-Stop-*XhoI* | CMV/pCDNA5 | Inactivation version of RNF5 |
| V5-UBB (expected MW : 10.3 kDa) | *AflII*-V5-*HindIII*-UBB-Stop-*XhoI* | CMV/pCDNA5 | For ubiquitination of vPOI |
| TOM20-pBirA(79aa-C term)-HA (expected MW : 45.0 kDa) | *KpnI*-TOM20-*BamHI*-linker-pBirA(79aa-C term)-HA-Stop-*NotI* | CMV/pCDNA5 | linker : SGGSGGSR TOM20(NM_014765.2) for applicating Contact-ID under ORF3a expression ref : Kwak et al., 2020 |
| Flag-pBirA(N term-78aa)-SEC61B (exepcted MW : 20.7 kDa) | *NotI*-Flag-pBirA(N term-78aa)-linker-*EcoRI*-SEC61B-Stop-*XhoI* | CMV/pCDNA3 | linker : GGASGGSGSGPVAT SEC61B (NM_006808) for applicating Contact-ID under ORF3a expression ref : Kwak et al., 2020 |
| ORF3a-twin strep (expected MW : 34.3 kDa) | *HindIII*-ORF3a-*BamHI*-twin strep-Stop-*XhoI* | CMV/pCDNA3.1 | |
| SEC61B-V5-TurboID (expected MW : 46.9 kDa) | *HindIII*-SEC61B-*BamHI-KpnI-ClaI*-V5-*NheI*-TurboID-Stop-*NotI-XhoI* | CMV/pCDNA5 | for identification of secreted proteins under ORF3a expression |

### Supplementary Table 2. ORF3a interactome : approved drug ID

| UNIPROT ID | Gene Name | Approved drug ID (drugbank.ca) : Drug name (Accession Number) |
|---|---|---|
| P05556 | ITGB1 | Antithymocyte immunoglobulin (DB00098) |
| O76082 | SLC22A5 | Levocarnitine (DB00583) |
| P53985 | SLC16A1 | Pyruvic acid (DB00119) |
| Q9UBM7 | DHCR7 | NADH (DB00157) |
| P29317 | EPHA2 | Dasatinib (DB01254), Regorafenib (DB08896), Fostamatinib (DB12010) |
| P30519 | HMOX2 | NADH (DB00157) |
| Q8WY07 | SLC7A3 | L-Lysine (DB00123), Fluciclovine (DB13146) |
| P19634 | SLC9A1 | Amiloride (DB00594) |
| Q16850 | CYP51A1 | Tioconazole (DB01007), Itraconazole (DB01167) |
| Q15738 | NSDHL | NADH (DB00157) |
| P38435 | GGCX | Coagulation factor VIIa (DB00036), Coagulation Factor IX (DB00100), Glutamic acid (DB00142), Menadione (DB00170), Phylloquinone (DB01022), Anisindione (DB01125), Kappadione (DB09332), Coagulation factor IX (DB13152) |
| P06241 | FYN | Dasatinib (DB01254), Fostamatinib (DB12010) |
| P05023 | ATP1A1 | Digoxin (DB00390), Acetyldigitoxin (DB00511), Hydroflumethiazide (DB00774), Etacrynic acid (DB00903), Trichlormethiazide (DB01021), Deslanoside (DB01078), Ouabain (DB01092), Diazoxide (DB01119), Bretylium (DB01158), Ciclopirox (DB01188), Bepridil (DB01244), Potassium cation (DB01345), Aluminium (DB01370), Magnesium cation (DB01378), Digitoxin (DB01396), Almitrine (DB01430), Magnesium acetate (DB13996), Potassium acetate (DB14498), Potassium sulfate (DB14499), Potassium (DB14500), Aluminium phosphate (DB14517), Aluminum acetate (DB14518) |
| P04035 | HMGCR | NADH (DB00157), Lovastatin (DB00227), Cerivastatin (DB00439), Simvastatin (DB00641), Atorvastatin (DB01076), Fluvastatin (DB01095), Rosuvastatin (DB01098), Pitavastatin (DB08860), Cannabidiol (DB09061) |
| P52429 | DGKE | Alpha-Tocopherol succinate (DB14001) |
| Q07954 | LRP1 | Antihemophilic factor (DB00025), Tenecteplase (DB00031), Coagulation Factor IX (DB00100), Coagulation factor IX (DB13152), Lonoctocog alfa (DB13998), Moroctocog alfa (DB13999) |
| P00533 | EGFR | Cetuximab (DB0000), Lidocaine (DB00281), Gefitinib (DB00317), Erlotinib (DB00530), Lapatinib (DB01259), Panitumumab (DB01269), Vandetanib (DB05294), Afatinib (DB08916), Osimertinib (DB09330), Necitumumab (DB09559), Foreskin keratinocyte (DB10772), Neratinib (DB11828), Dacomitinib (DB11963), Fostamatinib (DB12010), Brigatinib (DB12267), Zanubrutinib (DB15035) |
| P78348 | ASIC1 | Amiloride (DB00594) |
| P48651 | PTDSS1 | Phosphatidyl serine (DB00144) |
| O60658 | PDE8A | Caffeine (DB00201) |

### Supplementary Table 3. M interactome : approved drug ID

| UNIPROT ID | Gene Name | Approved drug ID (drugbank.ca) |
|---|---|---|
| P05556 | ITGB1 | Antithymocyte immunoglobulin (DB00098) |
| Q99808 | SLC29A1 | Troglitazone (DB00197), Ethanol (DB00898), Fostamatinib (DB12010) |
| P53985 | SLC16A1 | Pyruvic acid (DB00119) |
| O76082 | SLC22A5 | Levocarnitine (DB00583) |
| Q8WY07 | SLC7A3 | L-Lysine (DB00123), Fluciclovine (DB13146) |
| O95819 | MAP4K4 | Fostamatinib (DB12010) |
| P05023 | ATP1A1 | Digoxin (DB00390), Acetyldigitoxin (DB00511), Hydroflumethiazide (DB00774), Etacrynic acid (DB00903), Trichlormethiazide (DB01021), Deslanoside (DB01078), Ouabain (DB01092), Diazoxide (DB01119), Bretylium (DB01158), Ciclopirox (DB01188), Bepridil (DB01244), Potassium cation (DB01345), Aluminium (DB01370), Magnesium cation (DB01378), Digitoxin (DB01396), Almitrine (DB01430), Magnesium acetate (DB13996), Potassium acetate (DB14498), Potassium sulfate (DB14499), Potassium (DB14500), Aluminium phosphate (DB14517), Aluminum acetate (DB14518) |
| P29317 | EPHA2 | Dasatinib (DB01254), Regorafenib (DB08896), Fostamatinib (DB12010) |
| P19634 | SLC9A1 | Amiloride (DB00594) |
| P11166 | SLC2A1 | Carboxymethylcellulose (DB11059) |
| Q9UPY5 | SLC7A11 | Cystine (DB00138), Glutamic acid (DB00142), Riluzole (DB00740), Sulfasalazine (DB00795), Acetylcysteine (DB06151), Thimerosal (DB11590) |
| P30519 | HMOX2 | NADH (DB00157) |
| Q9UP95 | SLC12A4 | Potassium chloride (DB00761), Bumetanide (DB00887) |
| P52569 | SLC7A2 | L-Lysine (DB00123), Ornithine (DB00129) |
| Q07954 | LRP1 | Antihemophilic factor (DB00025), Tenecteplase (DB00031), Coagulation Factor IX (DB00100), Coagulation factor IX (DB13152), Lonoctocog alfa (DB13998), Moroctocog alfa (DB13999) |
| P49069 | CAMLG | Cyclosporine (DB00091) |
| Q16850 | CYP51A1 | Tioconazole (DB01007), Itraconazole (DB01167) |
| Q9Y666 | SLC12A7 | Potassium Chloride (DB00761) |
| P06241 | FYN | Dasatinib (DB01254), Fostamatinib (DB12010) |
| P78348 | ASIC1 | Amiloride (DB00594) |
| Q9UBM7 | DHCR7 | NADH (DB00157) |
| Q8TD43 | TRPM4 | Glyburide (DB01016) |
| P06213 | INSR | Insulin (DB00030), Insulin lispro (DB00046), Insulin glargine (DB00047), Insulin prok (DB00071), Mecasermin (DB01277), Insulin aspart (DB01306), Insulin detemir (DB01307), Insulin glulisine (DB01309), Chromic chloride (DB09129), Insulin degludec (DB09564), Fostamatinib (DB12010), Brigatinib (DB12267), Mecasermin rinfabate (DB14751) |
| P16435 | POR | Flavin adenine dinucleotide (DB03147), Flavin mononucleotide (DB03247) |
| P84077 | ARF1 | Glycerin (DB09462) |
| O60658 | PDE8A | Caffeine (DB00201) |
| P12931 | SRC | Dasatinib (DB01254), Citric acid (DB04272), Bosutinib (DB06616), Ponatinib (DB08901), Nintedanib (DB09079), Fostamatinib (DB12010) |

| | | |
|---|---|---|
| P04035 | HMGCR | NADH (DB00157), Lovastatin (DB00227), Cerivastatin (DB00439), Simvastatin (DB00641), Atorvastatin (DB01076), Fluvastatin (DB01095), Rosuvastatin (DB01098), Pitavastatin (DB08860), Cannabidiol (DB09061) |
| Q9HB21 | PLEKHA1 | Citric acid (DB04272) |
